# Rat superior colliculus encodes the transition between static and dynamic vision modes

**DOI:** 10.1101/2022.11.27.518086

**Authors:** Rita Gil, Mafalda Valente, Noam Shemesh

## Abstract

When visual stimuli are presented at a sufficiently high temporal frequency, visual perception shifts from the static to dynamic vision mode, thereby facilitating a continuity illusion which is key for correctly identifying continuous and moving objects and placing them in the context of the surrounding environment. However, how this continuity illusion is encoded along the entire visual pathway remains poorly understood, with disparate Flicker Fusion Frequency (FFF) thresholds measured at retinal, cortical, and behavioural levels. Here, we hypothesized that these disparities may suggest that other brain areas may be involved in encoding the shift from static to dynamic vision modes. We employ a comprehensive approach encompassing behavioural measurements, whole brain activation mapping with high fidelity functional MRI (fMRI), and local electrophysiological validation for studying the mechanisms underlying the shift from static to dynamic vision modes in the rat. Our behavioural measurements reported an FFF threshold proxy of 18±2 Hz. At the network level, functional MRI revealed that the superior colliculus (SC) exhibits marked signal transitions from positive to negative fMRI signal regimes at the behaviourally measured FFF threshold surrogates, with a strong linear correlation between fMRI signal and behaviour, while thalamic and cortical visual areas displayed a significantly poorer correlation with the behaviour. fMRI-driven neurometric curves approximated the behavioural psychometric curve in SC but not in the other visual areas. Electrophysiological recordings in SC suggested that these fMRI signals transitions arise from strong neural activation/suppression at low/high frequency regimes, respectively, and that a transition between these regimes occurs around the measured FFF threshold proxies. Lesions in V1 further reinforced that these transitions originate in SC. Combined, our data suggests a critical role for SC in encoding temporal frequency discriminations, in particular the shifts from the static to the dynamic vision modes.

## Introduction

The mammalian visual system^1–5^ has evolved ingenious ways for recognizing and extracting visual features that enable object perception^6,7^ and visual motion detection^8–11^, both essential for interacting with the external environment. The encoding of spatial resolution features along the entire visual pathway is well characterised, with most brain structures exhibiting topographical mappings^12–16^ that systematically represent the visual space. By contrast, how visual systems resolve luminance changes over time^17,18^ has yet to be explained on a systems level, with most studies focusing mainly on the retina^19–21^ and/or the visual cortex (VC)^22–24^.

A critical temporal phenomenon for visual encoding is the continuity illusion effect: when photons impinge on the retina, the visual pathway can operate in static vision mode – whereby every flash is encoded as a separate event promoting attention and novelty perception – or can shift to the dynamic vision mode, where flashing stimuli is “fused”, thereby producing a continuity illusion whereby light is perceived as a continuous and steady^17,25,26^. The Flicker Fusion Frequency (FFF) threshold defines the transition frequency from static to dynamic vision modes. Retinal cyto-organization strongly affects measured FFF thresholds^20,27,28^: For example, diurnal fast-moving animals such as birds possess high visual temporal resolution which enables the detection and processing of fast-moving stimuli, such as prey, obstacles, as well as maintaining formation when flying in flocks^25^. The FFF threshold also plays important roles in prey-predator interactions, for example, in camouflaging moving prey, or in detecting predators (a dynamically changing appearance can elicit a startle/fear response, giving prey an advantage to escape)^29^. Systemic medical conditions such as hepatic encephalopathy or eye disorders such as cataract or glaucoma can also strongly affect the FFF threshold and thus visual perception^17,25^.

Interestingly, FFF thresholds derived from behaviour^22,25,30–34^ and electrophysiological recordings^19–24^ are disparate. For example, hens do not appear to behaviourally perceive flicker frequencies above 75-87 Hz^25,35^, while their electroretinograms (ERGs) remained in phase with the flickering light well beyond 100 Hz^19,20^. Similar trends were observed in mice, where ERGs vs. behavioural reports of FFF thresholds were ∼30 Hz^18^ and ∼14 Hz^36^, respectively. Strikingly, electrophysiological recordings in the cortical end of the visual pathway disagreed both with behavioural- and with ERGs-derived FFF thresholds, further suggesting that behaviourally relevant FFF threshold encoding may occur elsewhere along the pathway.

Here, we combined behavioural measurements, network-level functional MRI (fMRI), and electrophysiological recordings, to investigate the network-level neural correlates of the transition from static to dynamic vision modes. We find that mechanisms boosting/suppressing neural activity in the Superior Colliculus (SC) are strongly associated with behavioural reports of static/dynamic vision modes, suggesting that the SC is a major junction for flicker fusion.

## Results

### Rats behaviourally report shifts from static to dynamic vision modes at 18± 2 Hz

We designed a simple psychophysical task to estimate the FFF threshold proxy in rats (**Figure 1)**. Rats were placed in a box with three ports and trained to report to one side port when the stimulus was continuous, and to the other when the light stimulus was flickering at 2 Hz (counterbalanced across animals; c.f. Methods). To ensure exposure to several flashes, even at low frequencies, rats were required to wait for 1 s before a tone signalled that a report was allowed. When animals chose the correct port, a water reward was delivered. Rats could perform this discrimination with an accuracy higher than 95% (**Figure S1**). The percentage of flicker port reports is shown in **Figure 1C**. For frequencies above approximately 20 Hz, rats predominantly chose the port rewarded to continuous stimuli. Notably, as the flicker frequency increases, the percentage of flicker port reports decreases, indicating that the animal tends to choose the continuous light port for higher frequencies. As the difficultly level for discriminating between individual flashes increases, the rats transition towards the dynamic vision mode: for the FFF threshold proxy calculation a sigmoid curve was fitted to the average animal response and the intercept at 0.5 (considered to be “chance level” as animals report equally to both ports) was taken. The calculated FFF threshold proxy was 18±2 Hz and the confidence interval was defined via a bootstrap method (c.f. Methods).

**Figure 1.**
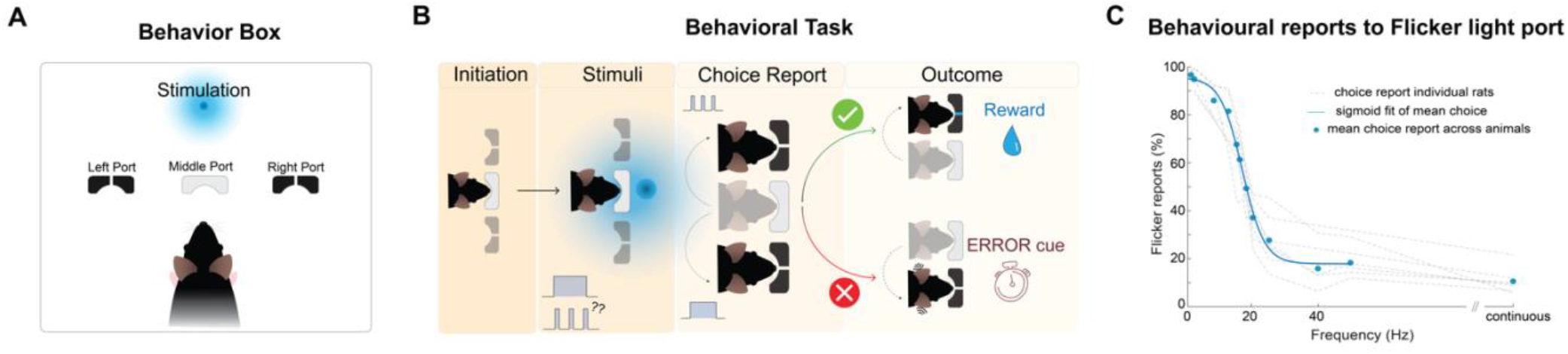
**(A) Behavioural results.** Water-deprived rats were placed in a dark box with three poking ports: a middle port to initiate trials and two lateral ports for continuous or flicker light reports. **(B) Schematic of the task:** Animals start each trial by poking in the central port. An overhead LED would turn on displaying either continuous light or flickering light at various frequencies. Rat were rewarded for poking to one side if the stimulus was continuous, and to the other side if the stimulus flickered. Incorrect responses triggered a noise burst and a time penalty. **(C) Percentage of reports to the flicker port.** Thin grey dashed lines reflect the performance of each individual animal (N=7) while blue circles correspond to the averaged individual performances. As the frequency increases the animal reports less often to the flickered port signalling a shift towards the dynamic vision mode. The calculated FFF threshold proxy at “chance level” is 18±2 Hz.

### Pathway-level fMRI reveals that SC signals, but not cortical or thalamic signals, tightly track behavioural reports

We then turned to investigate activity in the entire visual pathway of a separate group of rats (N=16) via functional-MRI (fMRI) experiments conducted at 9.4T with binocular flashing visual simulation (spatial resolution of ∼270×270 µm^2^ in-plane, 1.5 mm slice thickness and 1.5 s temporal resolution). Physiological conditions and MRI pulse sequence optimization were performed and shown in **Figure S2-5**. The stimulation paradigm, and LED positioning, relative to the animal’s eyes, is shown in **Figure 2A**. Functional activation t-maps for representative stimulation frequencies of 1, 15, 25 Hz and continuous light are shown in **Figure 2B**. At the lowest stimulation frequency, strong positive Blood-Oxygenation-Level-Dependent (BOLD) responses (PBRs) are observed in subcortical structures of the visual pathway (SC and thalamic lateral geniculate nucleus of the thalamus – LGN). Cortical areas exhibit somewhat weaker PBRs. As the stimulation frequency increased, gradual shifts were observed from PBRs to negative BOLD responses (NBRs), first in VC and then in SC. LGN responses remained positive for all frequencies, but t-values decreased with frequency. Temporal profiles in the anatomically defined regions of interest (ROIs) confirm the trends described above (**Figure 2C**), and further reveal sharp positive signals at the beginning and end of stimulation at the higher frequency stimuli in SC (hereafter referred to as onset and offset signals, respectively) flanking a lower “steady-state” fMRI response.

**Figure 2:**
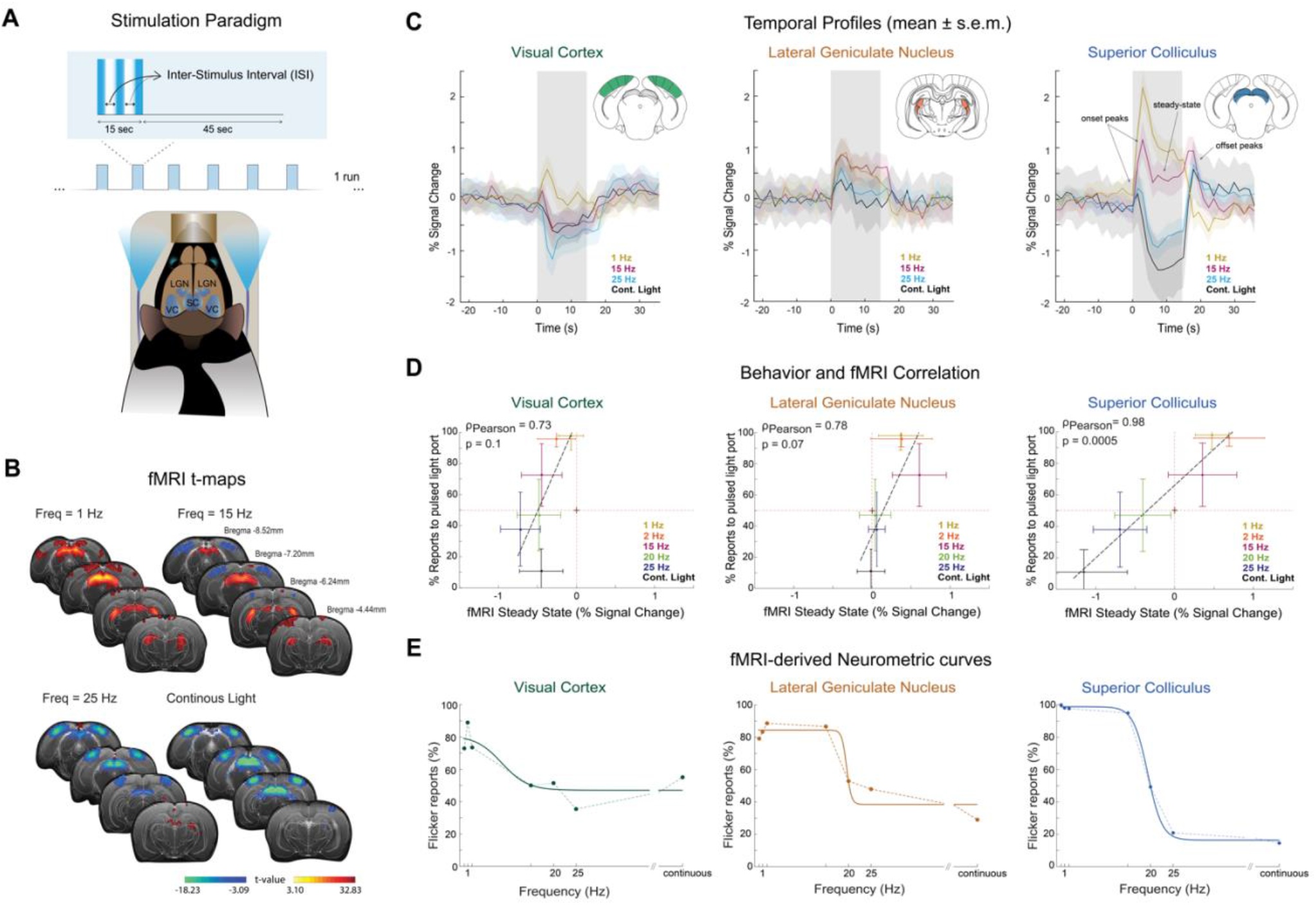
Whole pathway fMRI results. **(A) TOP: Stimulation paradigm used in the fMRI.** The stimulation paradigm consisted on 15 s stimulation at different frequencies followed by 45 s rest: **BOTTOM: Schematic animal’s position in MR bed. (B) TOP: Anatomical MRI image** with delineated ROIs and atlas overlapped. Different brain slices highlight visual pathway structures; **BOTTOM: fMRI t-maps for representative visual stimulation frequencies**. As the stimulation frequency increases, transitions from PBRs to NBRs are observed first in VC and subsequently in SC. **(C) Mean ± s.e.m. of fMRI signal temporal profiles across animals (N=16).** Fine structure appears in the fMRI responses. Onset and offset peaks, are evident in the SC profiles from 15 Hz onwards, along with a “steady-state” (black arrows); **(D) Correlation of behaviour reports with fMRI “steady-state” signals.** Coloured circles represent the average response of behavioural and fMRI sessions and error bars represent the standard deviation across runs/sessions. Only the SC shows a clear transition from PBR to NBR that correlates with the behaviourally measured FFF threshold surrogate; **(E) fMRI-derived neurometric curves** generated from individual trial data reveal that only SC tracks the psychometric curve obtained from the behavioural experiments, reporting a “chance level” threshold of 20.0±2.8 Hz.

To investigate the relationship between pathway-wide fMRI responses and behaviour results, we measured a larger stimulation frequency space and correlated the mean fMRI percent signal change across rats during the stimulation “steady-state” (c.f. Methods) with the mean probability of reporting flicker across animals (**Fig. 2D**). Interestingly, VC and LGN evidenced saturated responses: for frequencies above 15 Hz, the negative VC responses remained rather constant, and, in LGN, signals exhibited very small positive percent signal change above 20 Hz.

By contrast, a much broader dynamic range was observed in SC, where fMRI steady-state signal crossed zero between 15-20 Hz – close to the behaviourally measured FFF threshold surrogate. Furthermore, the SC is only structure that exhibits a highly significant correlation coefficient of *ρ*_*pearson*_ = 0.98 (P=0.0005, two-sampled *t*-test). Although this correlation is based on mean responses across 6 frequencies, it failed to reach significance in the VC and LGN with *ρ*_*pearson*_ = 0.73 (P=0.1, two-sampled *t*-test) and *ρ*_*pearson*_ = 0.78 (P=0.07, two-sampled *t*-test), respectively.

Finally, to quantify the single-trial discriminability of the fMRI responses as a function of flicker frequency, we computed fMRI-derived neurometric curves^75^ (**Figure 2E**) for each ROI. This was done by estimating the proportion of single-trial fMRI percent change responses (averaged across animals) above the mean value for the 20 Hz conditions, which is the closest frequency to the behaviorally derived FFF threshold surrogate (c.f. Methods for more details) For the SC ROI, but not for VC or LGN, the fMRI-derived neurometric function had a sigmoidal shape, which resembled the behavioural psychometric function, with a “chance-level” threshold of 20.0±2.8 Hz.

### Electrophysiology reveals that activation/suppression of neural activity underpin fMRI signal transitions in SC

As the SC showed the strongest behaviour-fMRI correlation, we targeted it for electrophysiological recordings (**Figure 3**). Fluorescence microscopy images (**Figure 3A**) validated the optimal angle of the silicon probe to record from the superficial layers of SC (SSC). **Figure 3B** and **3C** detail LFP traces (c.f. **Figure S6** for zoomed-in plots) and total spectral power over time between 1-50 Hz for representative stimulation conditions, respectively (c.f. **Figure S7** for remaining conditions). At 1Hz, individual flashes clearly elicited LFP periodic transients and strong power increases. At 15 Hz, sharp power increases are observed at the beginning and end of the stimulation block, while more modest power increases were observed during stimulation in the 15 Hz band (and some of its harmonics). By contrast, in the 25 Hz condition, periodic transients are much reduced while the sharp onset and offset signals are still observed (**Figures 3B, 3C and Figure S6**).

**Figure 3:**
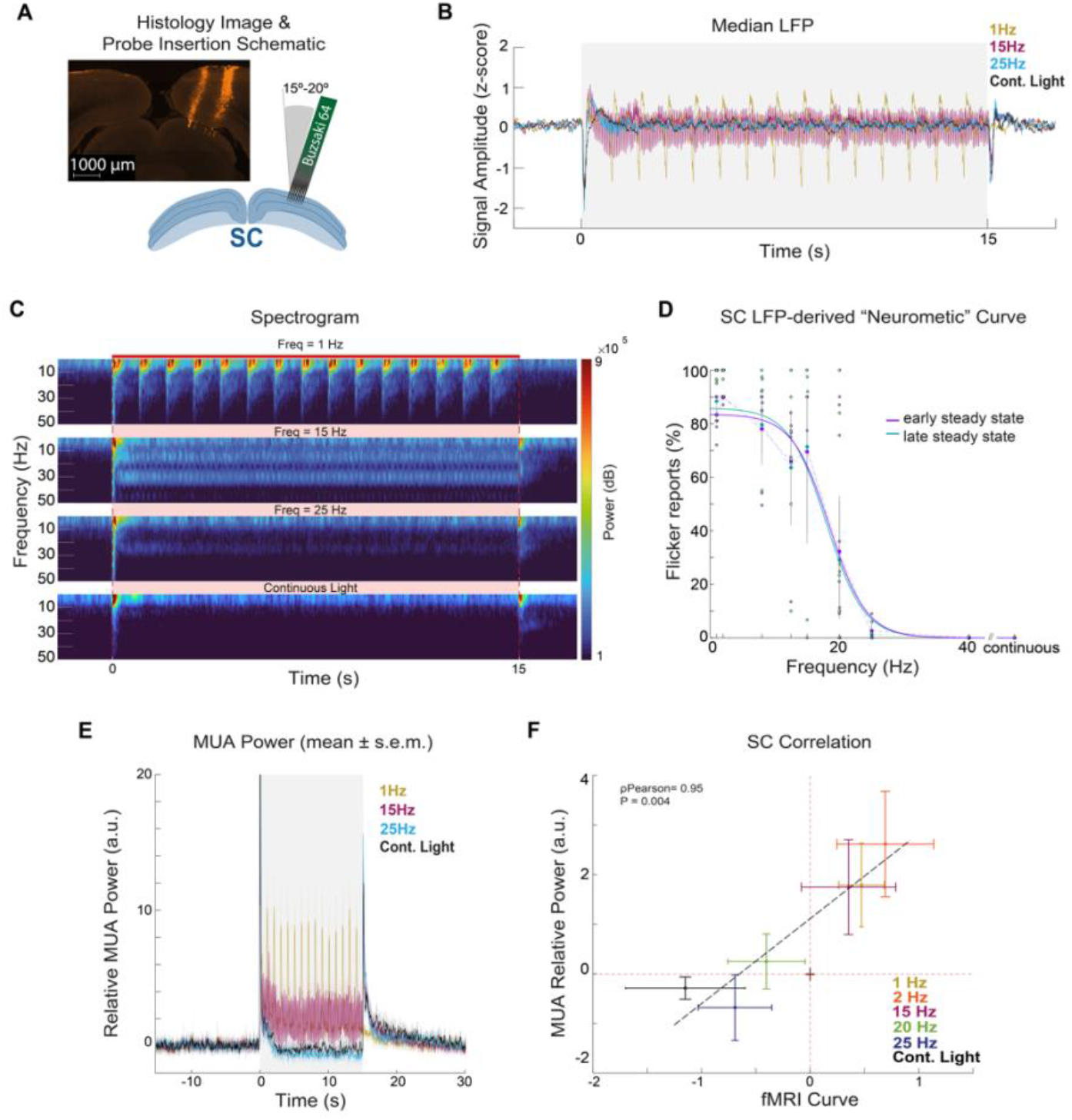
Electrophysiology results. **(A) Probe insertion schematic and fluorescence microscopy image; (B) Median LFP traces.** LFP traces show individual flash-induced LFP oscillations for the 1 Hz condition and onset and offset peaks for the higher frequencies; **(C) Spectrograms between 1-50 Hz for 1, 15, 25 Hz and continuous light stimulation regimes**; **(D) “Steady-state” LFP-driven neurometric curves.** The dots represent mean flicker reports for each animal while the error bars represent the 95% confidence interval. The fitted sigmoids reveal “chance-level” thresholds of 18.0±1.7 Hz and 17.7±2.4 Hz, for the early and late “steady-state” intervals, respectively. These values are in concordance with the ones obtain in the SC fMRI-driven “neurometric” curves and behavioural psychometric curve. **E) Mean ± s.e.m MUA relative power plots across animals**. These plots reveal a decreased power during stimulation as the frequency of stimulation increases. Offset signals become evident for the high frequencies. The two highest frequency regimes show power decreases during stimulation below baseline level; **(F) Correlation between “steady-state” MUA relative power and fMRI percent signal change.** Coloured circles represent the average response of electrophysiological and fMRI sessions and error bars represent the standard deviation across runs. A high correlation coefficient of ρ_Pearson_=0.95 (P=0.004, two sampled *t*-test) shows a tight relationship between the two measurements. NBRs at high stimulation frequencies correlate with strong MUA power reductions.

**Figure S6** further shows that individual flash-induced LFP power increases are present during the entire 1 Hz stimulation regime, while a “steady-state” is reached after ∼1-2 s of stimulation for 15 and 25 Hz. To investigate how discriminable was the “steady-state” periodicity in the transients evoked by stimuli of different frequencies in single trial, we computed LFP-derived neurometric curves for two different intervals beginning at 2 or 8 s after stimulation onset with a 5 s duration, respectively (**Figure 3D**). These curves were calculated by computing the fraction of trials in which the integral of the LFP power spectrum at the frequency of the stimulus used in that trial, was higher than the mean (across all the 20 Hz stimuli) of the integral of the LFP power spectrum at 20 Hz (c.f. Methods for more details). Both “steady-state” intervals revealed similar LFP-derived neurometric curves with chance-level thresholds similar to the ones obtained from the behavioural data and fMRI: 18.0±1.7 Hz and 17.7±2.4 Hz for early and late “steady-state” intervals, respectively.

When the MUA power along the different stimulating frequencies was computed (**Figure 3E)**, similar trends were observed. Interestingly, the “steady-state” for the two higher frequency conditions (25 Hz and continuous light) drops below baseline levels (suggesting inhibition), contrasting with the LFPs at this regime (**Figure S8**). To better explain the PBR to NBR transitions observed with fMRI in SC, we tested a broader range of visual stimulation frequencies (**Figure S8**). Higher stimulation frequencies clearly produced reductions in MUA power – suppression of neural activity – alongside larger NBRs.

**Figures 3F** investigates the relationships between fMRI time-courses and their MUA counterparts by showing the correlation between the mean “steady-state” signals of the two modalities (a similar analysis is shown for the LFP band in **Figure S9**). The two signals are highly correlated (*ρ*_*pearson*_ = 0.95 (P=0.004, two sampled *t*-test)). Stimuli below the chance-level threshold clearly induced fMRI and MUA positive signals, while stimuli above this threshold lead to a strong reduction of fMRI and MUAs signals.

The fMRI-BOLD signals are naturally delayed compared with their fast MUA counterparts due to the complex neurovascular coupling mechanisms^37–41^. **Figure S10** shows the 1 Hz and 25 Hz LFP and MUA curve convolved with an HRF. Interestingly, the convolved electrophysiological signals predict well the timing observed for the onset/offset fMRI signals timing (fast fMRI curves were acquired for this figure with a temporal resolution of 500 ms).

### V1 feedback acts as gain control for activation to suppression transitions in SC

Since SC receives cortical feedback from V1 (**Figure 4A**), we sought to investigate whether the transitions from PBR to NBR responses strongly depend on such feedback connections. To this end, V1 was bilaterally lesioned via ibotenic acid in 10 animals. Ex-vivo histological images (**Figure 4A**) and in-vivo structural MRI scans (**Figure 4B, top row)** confirmed the localization of the lesions in V1. Interestingly, secondary cortical regions now show enhanced PBRs at 1 Hz, while at higher frequencies, the NBRs observed in the sham were not apparent in the lesion group. Strikingly, the SC NBRs are attenuated in the lesion compared to the sham groups (**Figure 4B**). Signal time-courses comparing the two groups for three representative frequencies are show in **Figure 4C** for SC signals (cortical and thalamic responses in the lesion group are shown in **Figure S11**). Clearly, the V1 lesions produced stronger onset and offset peaks as well as weaker negative steady-state fMRI signals in the SC. Given that SC signals at lower frequencies are less affected by V1 lesions (**Figure 4C**), it may be deduced that V1 exerts a gain effect in SC beyond the FFF threshold, but is not necessary for producing suppression of activity in SC at higher flicker frequency. **Figure 4D** proposes a hypothesized mechanism for SC signals. We hypothesize that the SC response can be decomposed into three distinct parts: (i) novelty detection (onset/offset signals) overlapped with (ii) frequency discrimination perception – a constant effect of activation/suppression of SC activity modulated by stimulation frequency -and (iii) cortical gain control. The cortical gain modulates the SC responses as function of stimulation frequency – when frequencies are low, the cortex awaits further information, and when frequencies are high, it increases the suppression in SC to avoid instigation of the novelty detection. Other contributions within the visual pathway cannot of course be ruled out in participating in this interaction.

**Figure 4:**
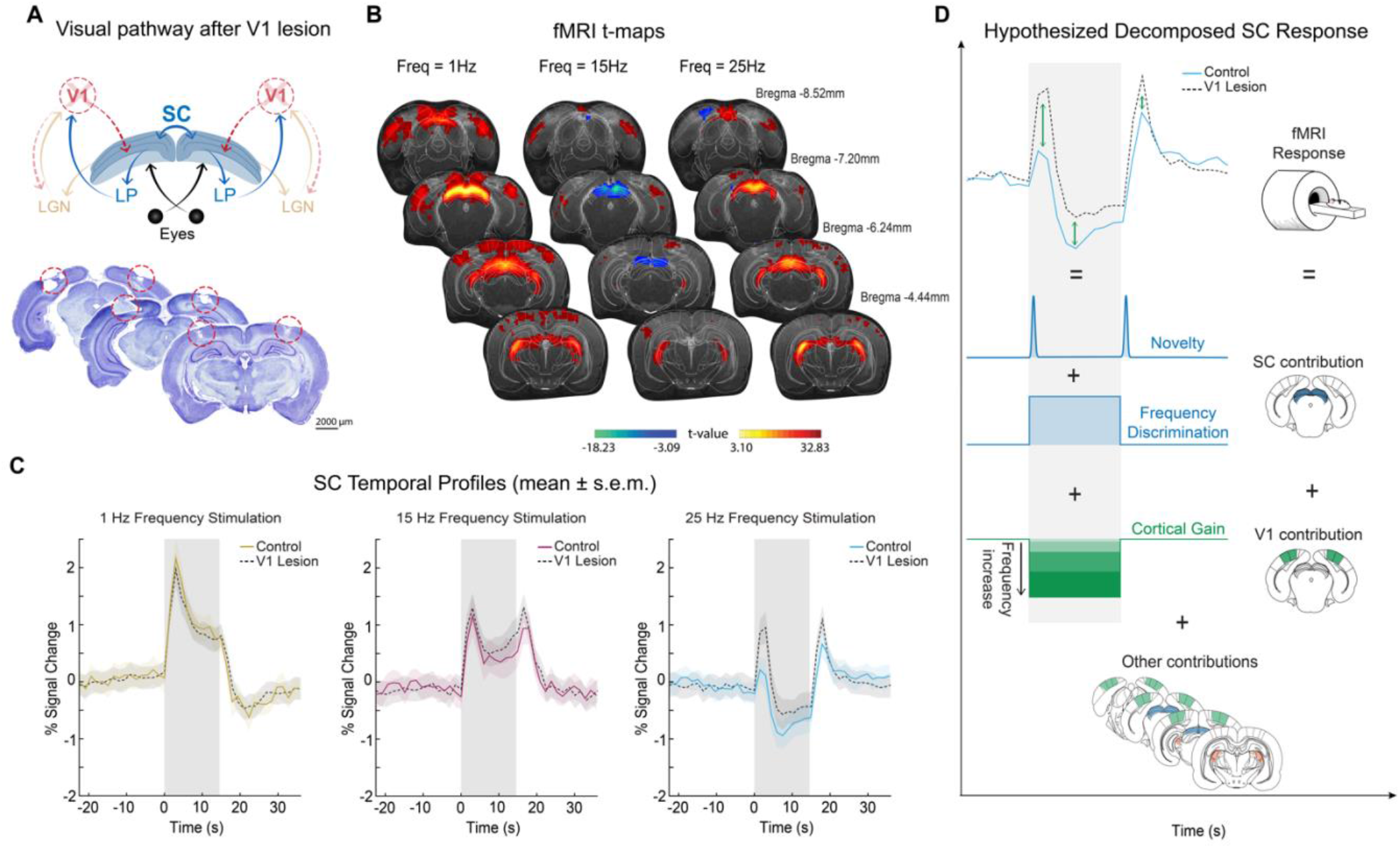
fMRI ibotenic acid lesions results. **(A) Schematic of visual pathway after V1 lesion** highlighting reduced cortical feedback **(TOP) and histological image (BOTTOM)** confirming the lack of brain tissue in the lesion site. **(B) TOP: Anatomical MRI image** with delineated ROIs and atlas overlapped; **BOTTOM: fMRI t-maps for representative frequencies**. As the frequency of stimulation increases, transitions from PBRs to NBRs appear in SC but not in VC. Secondary visual cortical areas show enhanced positive BOLD responses at 1 Hz compared to the control regime probably reflecting plasticity events that took place between the lesion induction and the fMRI experiments. **(C) Mean ± s.e.m. SC Temporal fMRI profiles across animals (N=10).** SC temporal profiles reveal marked positive to negative BOLD shifts and the onset and offset signals remain present after the lesion. Reduced amplitude of negative SC BOLD responses highlights the gain effect from V1 feedback projections; **(D) Hypothesized Decomposed SC Response.** An hypothesized mechanism is proposed based on the fMRI results. The SC response has at least 2 different contribution sources: a contribution within SC (novelty and a constant frequency discrimination perception) and a cortical contribution acting as gain control. Possible contributions from other structures within the visual pathway cannot be ruled out.

## Discussion

Investigating the brain’s ability to “fuse” sensorial inputs and induce complex mechanisms such as the continuity illusion (an analogue of which can also occur in the auditory domain^74^) is critical to better understand phenomena such as visual perception and its encoding along the visual pathway.

Studies focusing on the light entry point, the retina^19–21^, in species that are more reliant on vision such as monkeys^30,42^, dogs^32^, cats^22,33,43,44^ or birds^20,25,45^, initially proposed that FFF thresholds were solely dependent on retinal function and rod/cone composition^23,46,47^. However, behaviourally-derived FFF thresholds were always found to be lower than those derived from retinal electrophysiological recordings^20,25,35,36^. Hence, temporal resolution cannot be limited by the retina’s ability to resolve flickers, but rather reflects processing downstream in the visual pathway, where thresholds are likely modified at various stages^20,28^. Attempts to measure correlates of the visual continuity illusion at the higher order perception levels were carried out at the last neural information processing stage (VC)^22–24^, where the lowest FFF thresholds were reported in several species^48–50^. Thresholds for individual cortical cells varied across a broad range of frequencies and importantly, cortical FFF thresholds were lower than behaviourally-observed thresholds^23^. Hence, we hypothesized that the area most relevant for switching between dynamic and static vision modes must be located elsewhere, and applied a multi-level investigation to find it.

Our behavioural task was designed to avoid biasing animals towards one port and the reward system was carefully chosen to motivate animals to report their percepts about stimulus continuity (c.f. Behavioural task supplementary discussion for a more detailed discussion on the task’s contingencies). The FFF threshold surrogate measured here (18±2 Hz, **Figure 1C)** agrees well with prior literature (∼21 Hz in a different task^23,51^). Interestingly, the whole-pathway fMRI approach revealed that signals in SC – but not VC or LGN signals – bear a close relationship with the behavioural results: flicker reports above chance-level and associated PBRs indicated an activation of the SC, while below chance-level, strong NBRs were observed during the bulk of the stimulation period (apart from the onset/offset signals, vide-infra), which intensified as the stimulation frequency was increased. Subsequent electrophysiological recordings in SC lent further credence to the fMRI findings and revealed that suppression of MUA is responsible for the strong NBR responses.

A plausible mechanistic hypothesis for these signals in SC would suggest activation upon perceiving flickering light (PBRs) and suppression of neural activity (NBRs) at high frequencies where the light “fusion” mechanisms take place and the animals report to have entered in the dynamic vision mode. Taken together, our findings suggest that activation/suppression balances in the SC are important contributors for temporal frequency discrimination and potentially for entering the continuity illusion.

Anatomically, two visual sub-pathways co-exist: the extrageniculate pathway (including SC) and the geniculate pathway (including LGN). Experiments lesioning either one or the other revealed different roles in FFF threshold determination^22,33^: while the geniculate pathway mediates high flicker frequencies, the extrageniculate pathway mediates lower frequencies. The SC would behave as a lowpass filter limiting high frequency perception, in line with our findings that the SC plays an active role for temporal frequency discrimination. Experiments looking into the involvement of the VC in flicker discrimination revealed that, while humans became insensitive to any form of light stimulation following cortical ablation, lower-order species such as cats^52^ and albino rats^48^, were still able to discriminate flicker from steady light. This suggests subcortical “fusion” mechanisms in lower-order species. In this study, lesioning V1 revealed that this region is not necessary for generating the observed SC onset and offset peaks as well as the transition from activation to suppression, but rather, that it exerts a gain effect likely arising from cortical feedback into SC. This strengthens the conclusion that SC plays a major role in temporal frequency discrimination while also highlighting involvement of other structures along the visual pathway

While only few studies investigated SC in the context of the continuity illusion^22,33^, it was much more widely studied in the context of response habituation^53,54^ (RH), a phenomenon which, similarly to the continuity illusion effect, occurs at high stimulation frequencies. RH is expected to serve as a form of short-term memory for familiar versus novel information based on the dynamic adjustment of response thresholds. Studies^54–57^ have suggested a mechanism of feedback inhibition to be behind RH: the co-activation of excitatory and inhibitory neurons leads to a long-lasting inhibition blocking responses to subsequent stimulus presentation at high enough frequencies. The RH effect could be a contributor to the continuity illusion effect (probably among other mechanisms occurring long the visual pathway) and, if indeed the two phenomena are related, the continuity illusion effect in the SC would result, in part, from inhibitory processes - in line with the measured neuronal suppression at high stimulation frequencies both via fMRI and MUAs.

Another interesting finding in this study, is the ability of BOLD-fMRI to resolve the onset/offset peaks at the high frequency stimulation regimes. Onset peaks have previously been described with electrophysiology in the SC^58–60^ - at different stimulation frequencies, responses to the first flash remained similar and only subsequent responses appeared reduced in amplitude. However, to our knowledge, this is the first clear demonstration of offset peaks at high frequency regimes. Co-activation of excitatory and inhibitory neurons^57^ alone is unlikely to fully explain the observed offset signals, suggesting that the measured neuronal suppression occurring during the stimulation period likely reflect a more active process, and not solely the result of different lasting simultaneous effects or adaptation. One interpretation could be that when individual flashes are no longer perceptible, the entire stimulation period is “fused” an integrated as “one long flash” and the SC onset/offset peaks reflect brightness changes or novelty detections at the edges of stimulation – from dark to bright (onset peak) and from bright to dark (offset peak).

Finally, we believe our work provides important insight into the ongoing debate on the nature of negative BOLD signals and their underlying biological underpinnings^61–63^. The correlations between MUA and NBRs in this study strongly point to neuronal supression^63–72^ as the most probable scenario, and provide a system where the amplitude of the such signals can be modulated by a simple experimental variable – the stimulation frequency – which could serve as an experimental handle for future research into the mechanisms underlying the NBR neurovascular coupling.

Several limitations can be noted for this study. First, animals are awake during the behavioural task, while they are kept under light sedation during fMRI and electrophysiology data acquisition, which could potentially account for some slight differences in perceptual behavioural reports and neural/BOLD signals. Still the good agreement between the measurements suggests that this was not a critical effect here. Second, the animals made their behavioural reports after 1 s of stimulus presentation, while in the fMRI and electrophysiological sessions, stimuli lasted for 15 s. Our steady-state analyses (**Figure 3D**) were designed to take these differences into account, and indeed revealed that this is not a major confounding effect since, although the report takes place early on, the stimulus perception is maintained throughout the stimulation period. Furthermore, as shown in **Figure S10**, fMRI signals track MUA signals: the timing of NBRs onset/offset and steady state at 1 Hz and 25 Hz could be fully predicted from the MUA curves convolution with a conventional HRF – a good indication that “steady-state” fMRI signals can be used with good confidence as a proxy for the MUA recordings and consequently, also explain the behavioural reports.

## Conclusions

We investigated the shift between static and dynamic vision modes using a powerful multimodal approach encompassing behaviour, whole-pathway fMRI and electrophysiological recordings in SC, aiming to bridge the disparity between behaviourally-reported and retinal/cortical FFF thresholds. We find that FFF threshold proxies in SC, but not VC or LGN, agree with FFF surrogates from behaviour and electrophysiology. We have shown that in the SC, neural activity is highly suppressed above the measured FFF threshold and therefore activation/suppression balances in SC play a crucial role in encoding the transition from low to high frequency discrimination.

## Acknowledgments

This study was funded in part by the European Research Council (ERC) (agreement No. 679058), as well as by Fundação para a Ciência e Tecnologia (Portugal), project 275-FCT-PTDC/BBB-IMG/5132/2014. The authors acknowledge the vivarium of the Champalimaud Centre for the Unknown, a facility of CONGENTO which is a research infrastructure co-financed by Lisboa Regional Operational Programme (Lisboa 2020), under the PORTUGAL 2020 Partnership Agreement through the European Regional Development Fund (ERDF) and Fundação para a Ciência e Tecnologia (Portugal), project LISBOA-01-0145-FEDER-022170. MV thanks Fundação para a Ciência e Tecnologia for a PhD fellowship PD/BD/141560/2018. All authors would like to thank Dr. Alfonso Renart for suggestions and critical reading of the manuscript, Dr. Cristina Chavarrías for the implementation of the fMRI in the acquisition MRI sequences, Ms. Francisca F Fernandes for customized fMRI analysis MATLAB codes, Dr. Bruno Cruz, Dr. Tiago Monteiro, and Mr. Filipe Rodrigues for input regarding the behaviour experimental design, and Mr. Juan Castiñeiras and Mr. Tiago Costa for advice on data analysis. Finally, the authors would like to thank Ms. Teresa Serradas Duarte for insightful discussions.

## Methods

All animal care and experimental procedures were carried out according to the European Directive 2010/63 and pre-approved by the competent authorities, namely, the Champalimaud Animal Welfare Body and the Portuguese Direcção-Geral de Alimentação e Veterinária (DGAV).

In this study, N=53 adult Long-Evans rats were used: the behaviour task was performed in N=7 animals; fMRI was performed in N=32 animals – 16 animals for the different frequency stimulation acquisitions, 6 animals for the fast fMRI acquisitions and 10 animals for the ibotenic acid lesions acquisitions –; and electrophysiology was performed in N=14 animals. Animals weighted 370 ± 100 g with *ad libitum* access to food and water (except for the behavioural animals which were water deprived) and were under normal 12h/12h light/dark cycle. Data analysis was performed using Matlab(R) (The Mathworks, Natick, MA, USA, v2016a and v2018b).

### Animal Preparation

Animals were water-deprived for 3 days prior to the behavioural task and their weights monitored during the entire duration of the study. Weight loss was capped at 10% body weight while water-deprived.

Before the MRI and electrophysiological acquisitions, animals were anaesthetised with 5% isoflurane (Vetflurane, Virbac, France) for 2 min and were then moved to either the MRI bed or the stereotaxic set-up, respectively. Animals were later sedated with a medetomidine solution (1:10 dilution in saline of 1 mg/ml medetomidine solution - Vetpharma Animal Health, S.L., Barcelona, Spain) by injecting a subcutaneous bolus (bolus=0.05 mg/kg) while isoflurane dosage kept on being reduced. During fMRI experiments the medetomidine bolus was administered 5 min after the initial induction while during the electrophysiological recordings the bolus was administered after the surgery was done (∼40 min). Ten minutes after the bolus was administered a medetomidine constant infusion of 0.1 mg/kg/h, delivered via a syringe pump (GenieTouch, Kent Scientific, Torrington, Connecticut, USA), started. Isoflurane was graduallys reduced for the next 10 minutes, and the it was stopped and animals subsequently remained only under medetomidine sedation throughout the entire MRI and electrophysiological data acquisitions. To achieve efficient isoflurane washout, acquisitions were always started between 50-60 min after bolus injection. After the experiments, sedation was reverted by injecting the same amount of the initial bolus of a 5 mg/ml solution of atipamezole hydrochloride (Vetpharma Animal Health, S.L., Barcelona, Spain) diluted 1:10 in saline. At the end of the electrophysiological recordings, the animal was euthanized with a solution of sodium pentobarbital (delivered via intraperitoneal injection) and the brain was extracted and maintained in 4% PFA for a period of 12 to 24 h before slicing for further microscopy imaging.

### Behaviour

To investigate the biological underpinnings of temporal frequency discrimination, we measured FFF threshold surrogates behaviourally.

Water deprived animals (N=7, 14-20 sessions) were placed in a box with three ports and trained to associate each lateral port to either continuous or flicker light (**Figure 1A**). The central port was used for trial initiation and the animal had to poke in for ∼200 ms. After this time, the visual stimulus started: a blue LED on the top of the behaviour box turned on displaying either a continuous or a flickering light at different frequencies (keeping the pulse width of 10 ms and modulating the ISIs). The animal had to wait 1000 ms until a pure tone was played (with a frequency of 10 KHz), after which it could report its choice. The light would turn off only after the animal left the central port. The animals were trained to poke on one side for continuous light, and on the other side for flickering light (the contingency was counter balanced between animals). If the animal selected the correct port a water reward of 25 µl was delivered, otherwise a time penalty of 5000 ms initiated along with a burst of white noise to signal the mistake (**Figure 1B**).

During the training phase animals were presented with either true continuous light or a flicker frequency of 2 Hz. The initial fixation time and waiting period were set to 10 ms and 0 ms, respectively, and were incremented by 1 ms each time the animal completed a trial. After reaching a performance >80% with these two easy conditions, other frequencies were introduced and presented in a random manner.

During the task, 50% of trials consisted of continuous light while the other 50% consisted of a flickering light at different frequencies: 1, 2, 8, 12.5, 15, 16, 18, 20, 25, 40, and 50 Hz. This was done to prevent one of the pokes from delivering more rewards. Within the 50% of flicker trials, only 10% contained the higher frequencies (above 8 Hz): the so-called “probe trials” where the animal might enter the continuity illusion regime. These “probe trials” were rewarded in the flicker port. A supplementary discussion is presented in **Figure S1**.

### MRI Acquisitions

Functional magnetic resonance imaging (fMRI) data was acquired to observed Blood-Oxygenation-Level-Dependent (BOLD) contrast modulations induced by the different frequencies along the entire rat visual pathway. The experiments were conducted using a 9.4T Bruker Biospec MRI scanner (Bruker, Karlsruhe, Germany) equipped with an AVANCE III HD console including a gradient unit capable of producing pulsed field gradients of up to 660mT/m isotropically with a 120 μs rise time was used. An 86 mm quadrature coil was used for radio frequency transmission and a 4-element array cryoprobe^73^ (Bruker, Fallanden, Switzerland) was used for signal reception. The software running on this scanner was ParaVision^Ⓡ^ 6.0.1. During the entire duration of the fMRI experiments, animals breathed a mixture of 95% oxygen and 5% medical air. The animal’s temperature was monitored during the entire experiment (36.5 ± 1°C) with an optic fibre rectal temperature probe (SA Instruments, Inc., Stony Brook, New York, USA) and was regulated using a heating system consisting of circulating water. Respiratory rate was measured using a pillow sensor (SA Instruments Inc., Stony Brook, USA).

For correct slice placement along the visual pathway, an anatomical T_2_-weighted Rapid Acquisition with Refocused Echoes (RARE) sequence (**Figure 1B**) was used (TR/TE = 1600 / 36 ms, RARE factor = 8, Echo spacing = 9 ms; Averages = 3; FOV = 18 × 16. mm^2^, in-plane resolution = 168 × 150 μm^2^, slice thickness = 800 μm, t_acq_ = 1 min 3 s). Several control experiments were performed to determine the best conditions for data acquisition (**Figure S2-S5**). The functional MR imaging (**Figure 2**) was acquired using a Spin-Echo Echo-Planar Imaging (SE-EPI) sequence (TE/TR = 40 / 1500 ms, PFT = 1.5, FOV = 18 x 16.1 mm^2^, in-plane resolution = 269 x 268 μm^2^, slice thickness = 1.5 mm, t_acq_= 6 min 50 s).

### Electrophysiological Recordings

To understand the neural underpinnings of temporal frequency discrimination, MRI-targeted electrophysiological recordings were performed in the right superior colliculus (SC) inside a double-wall anechoic chamber, allowing for outside noise and light contamination reduction and acting as a Faraday cage.

Each recording lasted ∼5 h and both stimulation and sedation conditions were kept as similar to the MRI acquisitions as possible.

During surgery, the animal was kept under the effect of 3% isoflurane and the eyes were protected from light by covering them with an opaque gel (Bepanthen augen und nasensalbe). To avoid electrical noise, the animal’s temperature was kept within physiological parameters with the aid of an air-activated heating pad.

Following stereotaxic coordinates (using the 6^th^ Edition of Paxinos & Franklin’s rat brain atlas^74^) calculated from previous anatomical T_2_-weighted RARE MRI scans, a 2 x 3 mm craniotomy was opened in the skull above the right SC; and the dura was removed under a stereoscope using a small needle (BD MicrolanceTM 518 0.3 x 13 mm).

Extracellular recordings were performed using a 64 channel (8 shanks × 8 sites) silicon probe (Buzsaki64, NeuroNexus, Ann Arbor, MI). The probe was connected to an analogue 64-channel headstage (Intan) and positioned with a manual micromanipulator at an angle with the vertical between 15°-20°, for better alignment with the SC and in order to avoid blood vessels. Prior to brain insertion, the probe was stained with DiI (Vybrant™ DiI Cell-Labelling Solution) for post-hoc confirmation of the recording location with microscopy imaging (**Figure 3A**). The skull cavity was kept moist with saline during this entire process.

The data was digitised at 16 bit and stored for posterior offline processing using an Open Ephys acquisition board (Open Ephys) at a 30 kHz sampling rate.

### Visual Set-Up and Paradigm for fMRI and Electrophysiological Acquisitions

The eyes of the animal were hydrated during the acquisition with an ophthalmic gel (Visidic gel Bausch + Lomb) and a bifurcated optic fibre connected to a blue LED (*λ*=470nm and I=8.1×10^−1^ W/m^2^) was placed horizontally in front of each eye of the animal for binocular visual stimulation (**Figure 2A**).

For the MRI acquisitions, the blue LED was connected to an Arduino MEGA260 receiving triggers from the MRI scanner and used to generate square pulses of light. For the electrophysiological acquisitions, the blue LED was connected to an I/O board built in-house and controlled through Matlab 2016a.

Several stimulation frequencies were used - 1, 2, 15, 20, 25 Hz and continuous light. For the flickering stimuli the flash duration was kept constant at 10 ms in order to modulate inter-stimulus intervals (ISIs).

The stimulation paradigm consisted of a 15 s stimulation period interleaved with 45 s rest periods (**Figure 1A**). Both the fMRI and electrophysiological acquisitions consisted of an initial resting period of 45 s followed by 6 and 10 cycles of the experimental paradigm, respectively.

### Ibotenic Acid Lesions

To investigate the effect of cortical feedback projections in SC signal modulations, animals (N=10) were injected bilaterally with an ibotenic acid^75^ solution (excitotoxic agent, 1mg/100uL) in the primary visual cortex (V1). The acid was injected using a Nanojet II (Drummond Scientific Company). Animals were anesthetized with isoflurane (anesthesia induced at 5% concentration and maintenance below 3%), and a scalpel incision was made along the midline of the skull, the skin retracted and the soft tissue cleaned from the skull with a blunt tool.

Coordinates for the craniotomies and amounts required for the injections were determined for each individual animal based on T_2_-weighted anatomical images acquired before the surgery. We injected in 5 different AP coordinates; 1 injection site for the first AP coordinate (2 pulses of injection); 2 for the following (4 pulses of injection each), for full coverage of V1. Each injection pulse was administered at a rate of 23 nL/s and 2-3 s between pulses and each pulse consisted on 32nL. Waiting time before removing the injection pipette after the last pulse was of 10 min. The craniotomies were then covered with Kwik-Cast™ and the scalp sutured.

## Data Analysis

### Behaviour Analysis

We analysed the behavioural task as a detection task^76,77^, in which the rat had to detect whether a flashing stimulus or continuous light were presented. The percentage of responses to the flicker light port was calculated and, to estimate the FFF threshold surrogate, a sigmoid function was fitted to the averaged animal response and the intercept at 0.5 (“chance level”) was taken. For the standard deviation calculation, performances were separated by individual sessions and a bootstrap method was applied by resampling with replacement (50 iterations). For the correlations with fMRI results the average performance of all sessions was computed and the standard deviation across sessions is shown in the error bars.

### fMRI Analysis

For the MRI data, a general linear model (GLM) analysis was conducted along with a region of interest (ROI) analysis to investigate temporal dynamics of activation profiles.

**GLM Analysis:** Pre-processing steps included manual outlier removal (a spline interpolation was made taking the entire time course), slice-timing correction (using a sinc-interpolation) followed by head motion correction (using mutual information). Data was afterwards co-registered to the T_2_-weighted anatomical images, normalised to a reference animal and smoothed using 3D Gaussian isotropic kernel with full width half-maximum corresponding to 1 voxel (0.268 mm).

The stimulation paradigm was convolved with an HRF peaking at 1 sec. A one-tailed voxelwise t-test was performed, tested for a minimum significance level of 0.001 with a minimum cluster size of 20 voxels and corrected for multiple comparison using a cluster false discovery rate test (FDR).

**ROI Analysis:** For the ROI analysis, the 6^th^ Edition of Paxinos & Franklin’s rat brain atlas^74^ served as guidance for the manual ROI delineation (**Figure 2B**). The individual time-courses were detrended with a 5^th^ degree polynomial fit to the resting periods in order to remove low frequency trends and were then converted into percent signal change relative to baseline. For each run, the six individual cycles were separated and averaged across all animals to obtain the averaged response within each ROI (along with the standard error of the mean).

For the “steady-state” time portion calculation the averaged fMRI response per stimulation frequency for each ROI was resampled to a 4 Hz sampling frequency. A comparison was then performed between two consecutive sliding windows comprised of 5 time points (sharing one time point). The “steady-state” was assumed to be reached when the difference between medians was less than 0.04 % signal change. The “steady-state” percent signal change for each run for different animals was then averaged and the averaged baseline value was subtracted in order to correct for any remaining baseline fluctuations. The mean fMRI percent signal change during “steady-state” across animals was then correlated with the mean percentage of reports to the flicker port (**Figure 2D**) and with the mean MUA relative “steady-state” powers (**Figure 3F**). The average across runs/sessions is represent in the correlation plots by the error bars.

fMRI-driven “neurometric” curves were calculated using the “steady-state” portion of fMRI signals from individual trials in order to quantify the single-trial discriminability of the fMRI responses as a function of flicker frequency. A threshold based on the behaviourally derived FFF threshold surrogate (20 Hz) was chosen. For each trial (cycle) we estimated the proportion of fMRI percent change responses (averaged across animals) which were above the mean value for the 20 Hz condition. A sigmoid function was fitted to the results (as done for the behavioural data) and the “chance level” threshold (FFF threshold proxy) was considered to be the 0.5 of the fit as in the behaviour experiments.

### Electrophysiology Analysis

Electrophysiological data was band-passed with a notch filter at 50, 100 and 150 Hz to remove power line noise and detrended with a linear fit to the entire run.

For each individual run, time-courses were divided into individual cycles and a weighted average across channels was performed in order to obtain the averaged channel cycles for each run based on the integral during stimulation period of the absolute signal for the lowest stimulation frequency of the group.

The power spectral density (PSD) was calculated using Welch’s estimate. A hamming sliding window with 25% overlap was used; the final temporal resolution was set to be 50 ms. We further filtered signals in the local field potential (LFP) frequency band (1-150 Hz) and computed the averaged signal for all animals for the different stimulation frequencies (**Figure 3B and S6**).

LFP-derived “neurometric” curves (averaged across rats) were computed to investigate how discriminable was the “steady-state” periodicity in the transients evoked by stimuli of different frequencies in single trials. For the calculation of such curves the power spectrum of each normalized LFP “steady-state” window signal per trial was computed. We further subtracted to these power spectra the averaged run power spectrum corresponding to a non-stimulation resting period (with the same duration) in order to remove contributions that did not come solely from the stimulation itself. As metric of comparison for each trial we used the peak integral around the stimulation frequency used in that trial and computed the fraction of trials in which this metric was higher than the mean, across all 20 Hz trials, of the peak integral of the LFP power spectrum at 20 Hz. A sigmoid function was then fitted to the results and the “chance level” threshold was calculated (0.5 of the fit). We computed the LFP-derived “neurometric” curves for two different time intervals: an early and late “steady-state” intervals defined as 2-7 s and 8-13 s after stimulation started, respectively.

For each run, the PSD integral in the desired frequency bands (LFP and multi-unit activity - MUA: 300-3000 Hz) was divided by each band width, and z-scored before averaging all runs from different animals (**Figure 3E** and **Figure S8**). The “steady-state” power was considered to be the average of power during stimulation after the first second until the end of the stimulation period. The mean “steady-state” power for MUAs and LFPs across animals, Figure 3F and S8 respectively, was correlated with the mean fMRI “steady-state” signals across animals to further investigate the relationship of these two data modalities. The error bars shown in these correlation plots represent the averaged of each run across animals. The lowest tested frequency (0.25 Hz) was excluded in this correlation since at such low stimulation frequency, the four individual flashes that occur during the stimulation period are distant enough to be perceived as four distinct stimuli and signals never reach a “steady-state”. For the 1 Hz and 2 Hz condition the individual flash-induced MUA and LFP power increases are still clearly separated but we assumed the mean power increase to be the “steady-state” power.

## Supplementary Information

### Supplementary Results and Figures

#### Behaviour

##### Behaviour movie

The file “**SI_behaviourMovie.mp4**” depicts a representative part of the behaviour sessions. The animal starts new trials by poking on the central port and then choosing the side ports depending on the frequency displayed on the above head light (on the selected session the continuous and flickering lights would be rewarded in the right and left ports, respectively).

##### Behaviour Discussion

Most studies investigating the continuity illusion have used a GO/NOGO task paradigm^30,36,51^ where animals were trained to respond in one port in order to report one of two conditions, flickering or continuous stimulus and to withhold any response in the case of the opposite condition. In the present task we went one step further and allowed animals to freely choose between two ports where each port corresponded to one light condition.

In the context of continuity illusion studies, behavioural tasks have been designed and constrained to approximate measures to FFF thresholds; however, to completely rule out the possibility of animals simply comparing low vs high frequencies instead of flicker vs continuous light would require complementary and more complex behavioural experiments than employed so far. Since results of such experiments may also depend on task design^36^, we focused on ensuring that our task would (i) avoid biasing the animals towards one side port; (ii) ensure that 50% of the trials would deliver continuous light; (iii) reward “probe trials” in the flicker port to prevent the low vs high frequency possible comparison from taking place; and (iv) limit presentation of frequencies above 8 Hz (“probe trials”) to only 10% of the flicker trials so that animals would not perceive the reward in the continuous stimulus as uncertain.

Despite these efforts, the observed behavioural percentage of reports to the flicker port for the continuous stimulus conditions reaches 10.7% instead of the expected values closer to 0%. This decrease in performance is not present for the easy flicker conditions, where reports are above ∼95%. This increase in “flickering” reports for the continuous light condition could represent a limitation at two different levels: (1) a limitation in the animals’ ability to discriminate the “true continuous”; or (2) a limitation in the design of the task. We theorize that the observed increase in “flickering” reports for the continuous light might be due to a limitation in the design of the task and not animal related. In line with this, during the initial phase of training, when animals are presented with only 2 Hz and continuous light, they reach performance levels of 98% and 97%, respectively. If the animals’ ability to discriminate “true continuous” light was indeed impaired then the percentage of “flickering” reports would already manifest in the initial training phase. This hints at the possibility that showing “probe” trials (that can potentially be perceived as continuous but were rewarded in the flicker port) for 10% of flickering trials might still have been excessive; and/or that the rewarding system used was suboptimal and negatively influenced the animals’ behaviour.

When calculating the FFF threshold proxies for such a complex behaviour, one could argue that other parameters, besides the animals’ performance, might have been taken into consideration. In **Figure S1** we show percentage of aborted trials (**Figure S1B**) as well as movement (**Figure S1C**) and reaction times (**Figure S1D**) for the different presented frequencies of stimulation. Supporting the measured FFF threshold proxy, animals’ show an increased percentage of aborted trials (by not waiting the required 1000 ms) in the vicinity of the calculated threshold (highlighted in red) and a small decrease in reaction and movement times after the calculated threshold (highlighted in red). However, it is unclear how other parameters could be easily included and we consider this to be out of the scope of this paper.

**Figure S1:**
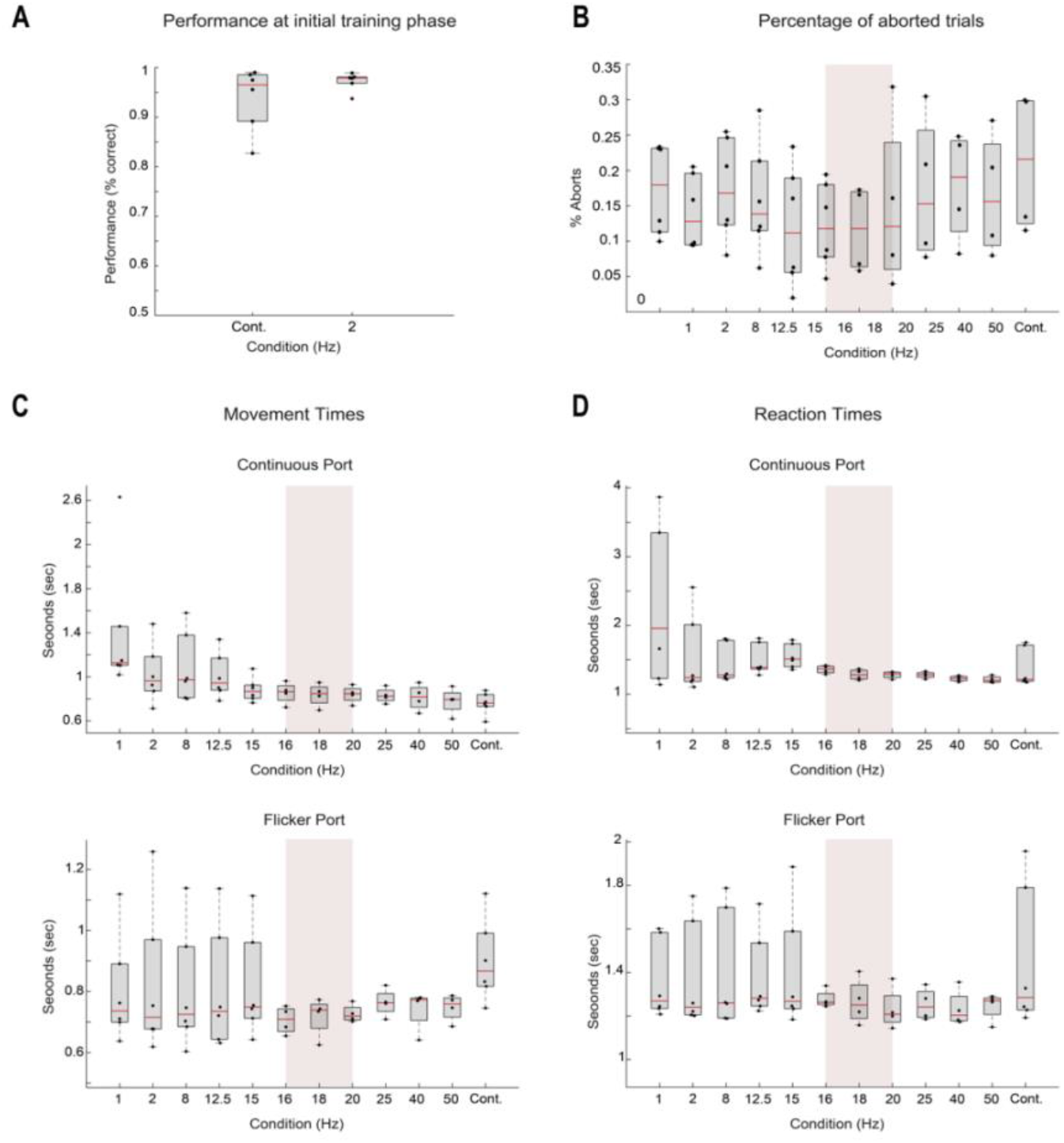
Behaviour Results. **(A) Performance at initial training phase.** Average performance of the animals for the initial training phase that included only the 2 Hz and “true continuous” stimuli. The values are shown for each of the presented frequencies and the mean is shown over animals with N=7; **(B) Percentage of aborted trials.** Percentage of trials aborted due to the animals attempting to respond before the mandatory minimum reaction time (1 second) reaching its end and the pure tone signalling the response period being played. The values are shown for each of the presented frequencies and the mean is shown over animals with N=7; **(C) Movement times**. Average over animals (N=7) of the movement times - time it takes the animal to reach the response port once it leaves the central port after the 1 second minimum-reaction time has elapsed - registered for the different frequencies presented. These are organized according to the report port for the trial, continuous port (top) and flickering port. The red-shaded area marks the calculated FFF ±2 Hz. **(D) Reaction times**. Average over animals (N=7) of the reaction times - time the animal is exposed to the stimulus, 1 second mandatory time plus the time the animal choses to linger until ready to respond - registered for the different frequencies presented. These are organized according to the report port for the trial, continuous port (**top**) and flickering port (**bottom**). The red-shaded area marks the calculated FFF threshold: 18±2 Hz.

#### Functional MRI

##### Oxygenation Percentage influence on fMRI responses

Different oxygenation levels (medical air - 21% O_2_; oxygen enriched air - 28% O_2_; and hyperoxia - 95% O_2_) were tested for two stimulation frequencies (2 Hz and 15 Hz) to investigate the sensitivity to measured positive and negative fMRI responses (**Figure S2**). Images were acquired with a SE-EPI sequence (TE/TR=43/1500 ms; FOV=18×16.1 mm; FA= 62°, resolution = 269×268 μm^2^; slice thickness=1.5 mm). Results reveal that, for both frequency regimes, the hyperoxia oxygenation state (95% O_2_) is the best regime to maximize fMRI responses, both positive and negative, along the rat visual pathway.

**Figure S2:**
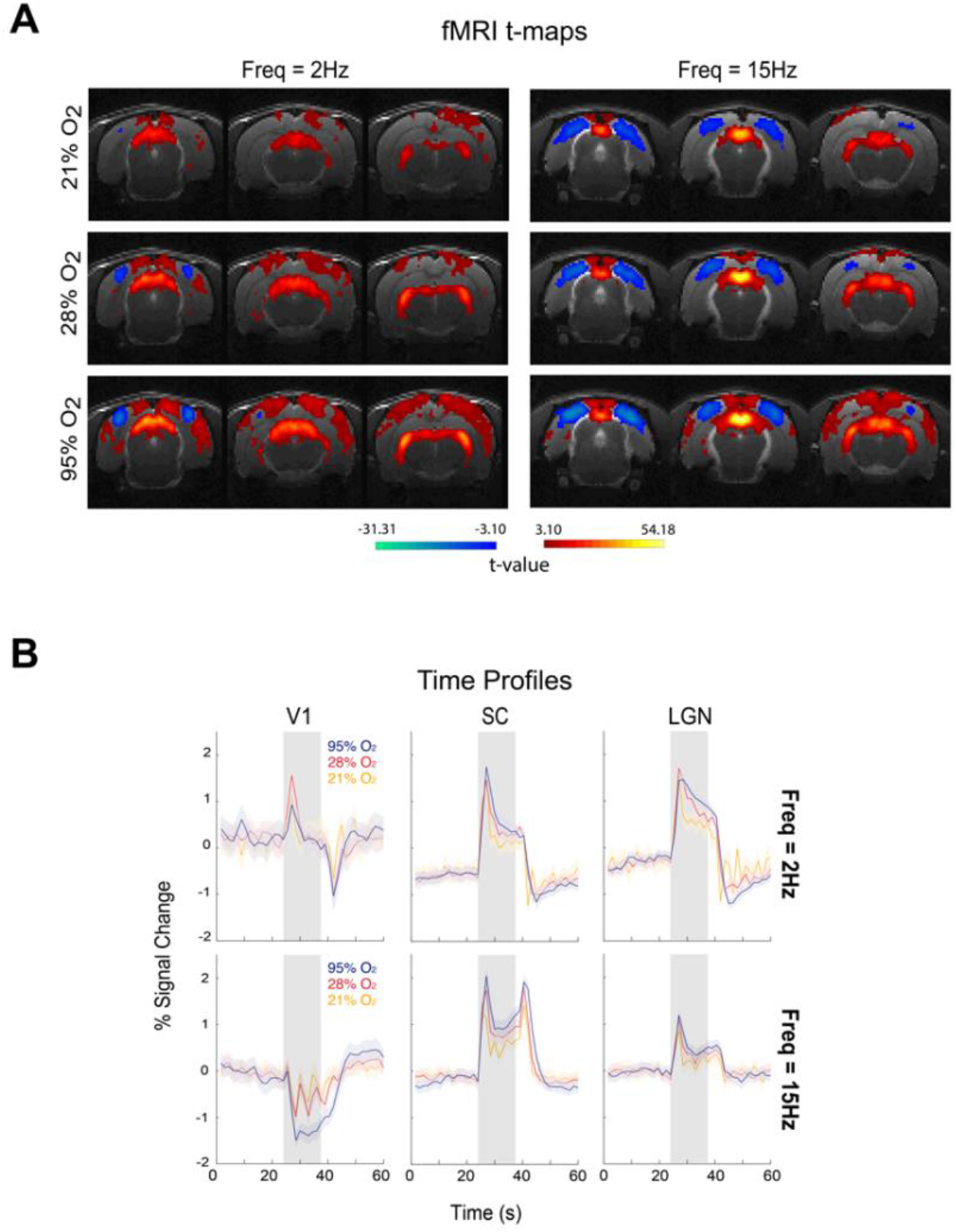
fMRI signal modulation with different percentages of oxygen. **(A) fMRI t-maps** for 23, 28 and 95% O_2_ (p-value=0.001, cluster size=20 voxels). Bilateral fMRI responses along the main structures of the visual pathway are observed for both stimulation frequencies. Negative cortical fMRI responses along with positive subcortical fMRI responses are emphasized as the percentage of oxygen increases. For the 95% O_2_ regime (hyperoxia) negative fMRI responses are observed in the V1M sub-regions (the monocular sub-region of the primary visual cortex) surrounded by positive responses in the rest of the primary visual cortex and secondary visual cortices. (**B) fMRI temporal profiles** for the different structures. An amplification of both positive and negative fMRI responses is confirmed for the hyperoxia condition. The amplification is mostly observed in cortical regions and more modest in subcortical regions. Legend: 21% O_2_ (medical air)– yellow; 28% O_2_ (oxygen enriched air) – orange; 95% O_2_ (hyperoxia) – blue.

##### Spin Echo vs. Gradient Echo fMRI acquisitions

To investigate if fMRI signal modulations observed at different stimulation frequency were specific to the chosen sequence, spin echo (SE) EPI, we tested several stimulation conditions with a gradient echo (GE) EPI sequence with similar parameters (TE/TR=15/1500 ms; FOV=18×16.1 mm; FA= 62°, resolution = 269×268 μm^2^; slice thickness=1.5 mm) as in the SE-EPI sequence.

Results shown in **Figure S3** reveal similar signal modulations observed in GE-EPI fMRI as the ones observed in SE-EPI acquisitions. Negative cortical responses appear first at 15 Hz stimulation frequency however, these appear more modest than the ones observed with SE-EPI acquisitions due to GE-EPI specific artifacts in these regions. SC responses show similar shifts from positive to negative responses as the stimulation frequency increases (at 25 Hz strong negative SC responses are observed). Due to enhanced specificity, the SE-EPI sequence was chosen for this study.

**Figure S3:**
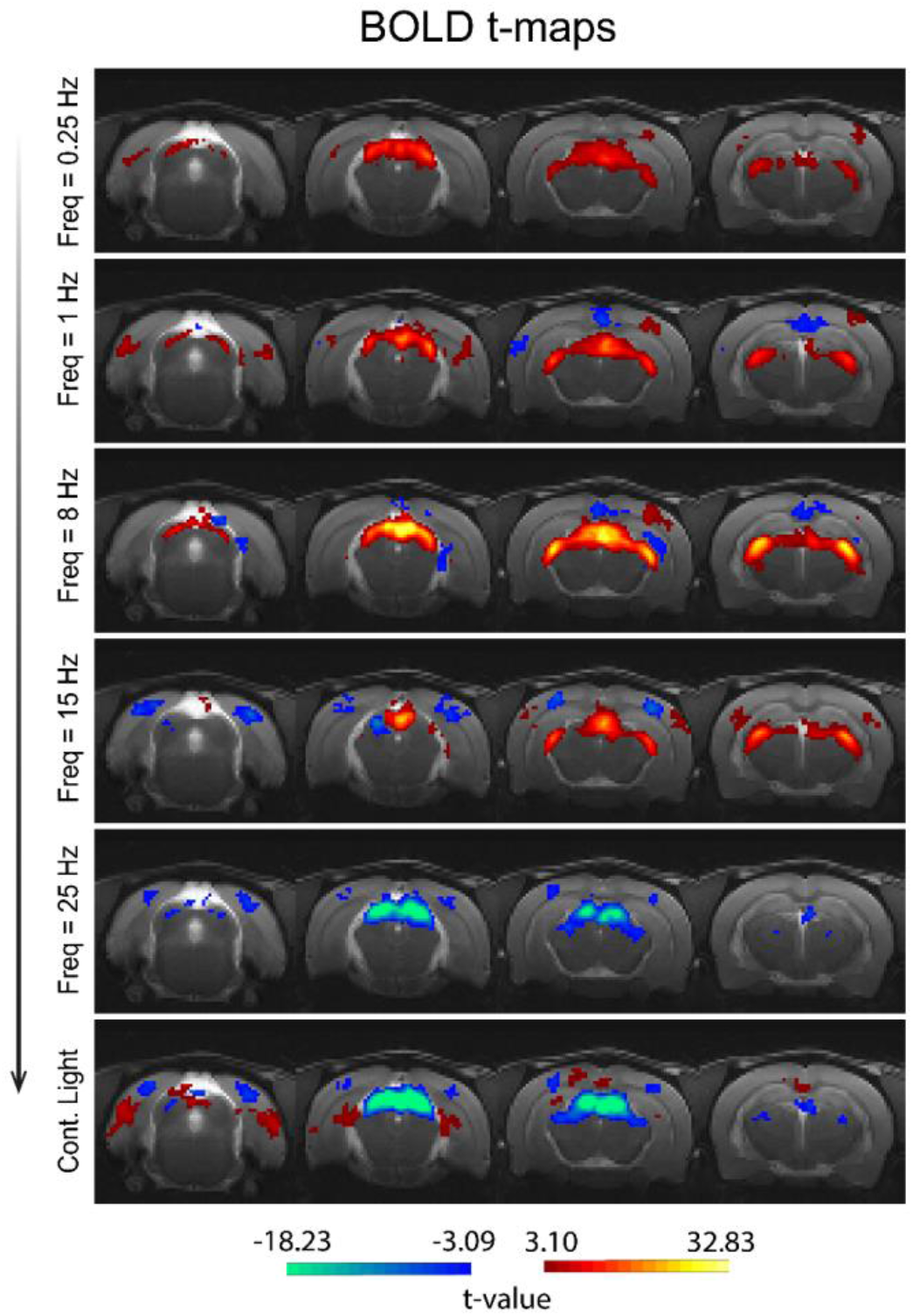
**GE-EPI fMRI t-maps along different stimulation frequencies** (cluster FDR corrected, p-value=0.001, cluster size=20 voxels). Bilateral fMRI responses are observed at the main visual pathway structures: SC, LGN and VC. Positive cortical responses at low stimulation frequencies are barely noticeable while stronger negative responses at frequencies higher than 15 Hz can be observed as in SE-EPI acquisitions. Subcortical responses also present signal modulations with strong negative responses in SC for frequencies higher than 25 Hz and reduced. LGN responses also appear modulated.

##### Stimulation Parameters influence on measured fMRI responses

From previous results we know that different frequencies with a constant flash duration of 10 ms (and consequently different ISIs) induce fMRI signal modulations. To further investigate the stimulation parameter (ISI vs flash duration) that has major influence in such signal modulations we tested fMRI signal modulations keeping the same ISIs but modulating the flash duration from 10 ms to 1 s (**Figure S4**). These acquisitions were performed using the chosen SE-EPI sequence: TE/TR=40/1500ms; FOV=18×16.1mm; resolution = 269×268μm^2^; slice thickness=1.5mm.

**Figure S4:**
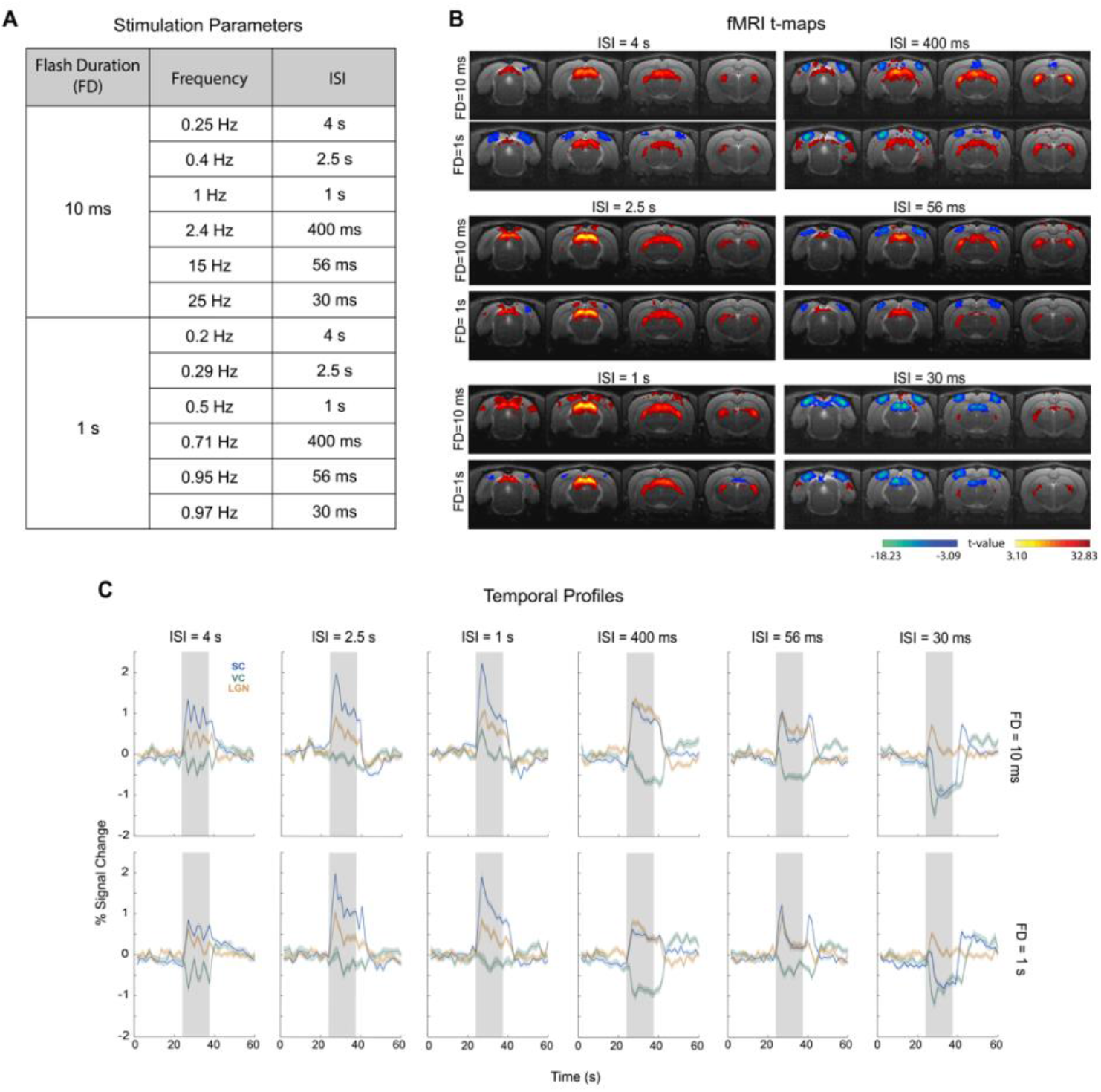
Effects of stimulation parameters on fMRI response modulations along the visual pathway. **(A) Stimulation Parameters** for the different tested conditions. Two flash durations (FD) were tested, 10 ms and 1 s, while keeping similar ISIs. **(B) fMRI t-maps** (p-value=0.001, cluster size=8 voxels, FDR cluster correction). Maps appear similar for both flash durations along the tested ISIs with increased cortical and subcortical negative fMRI signals at high frequencies; **(C) fMRI Temporal Profiles.** To further investigate the how stimulation parameters affected fMRI signal modulations we plot the temporal profiles of signals with different FDs. These plots show similar curves for the three different ROIs (superior colliculus, SC, lateral geniculate thalamic nucleus, LGN, and visual cortex, VC) for the two different FDs. Modulations appear similar as the ISIs decrease. These results reveal that the ISI is the major factor inducing such changes.

Results show that different flash durations originate similar results revealing that the change in ISIs is the main factor inducing fMRI signal modulations.

##### Average time-courses for different stimulation frequencies

Time courses from the chosen SE-EPI sequence are shown in **Figure S5** for the different drawn ROIs along the visual pathway and different stimulation frequencies.

**Figure S5:**
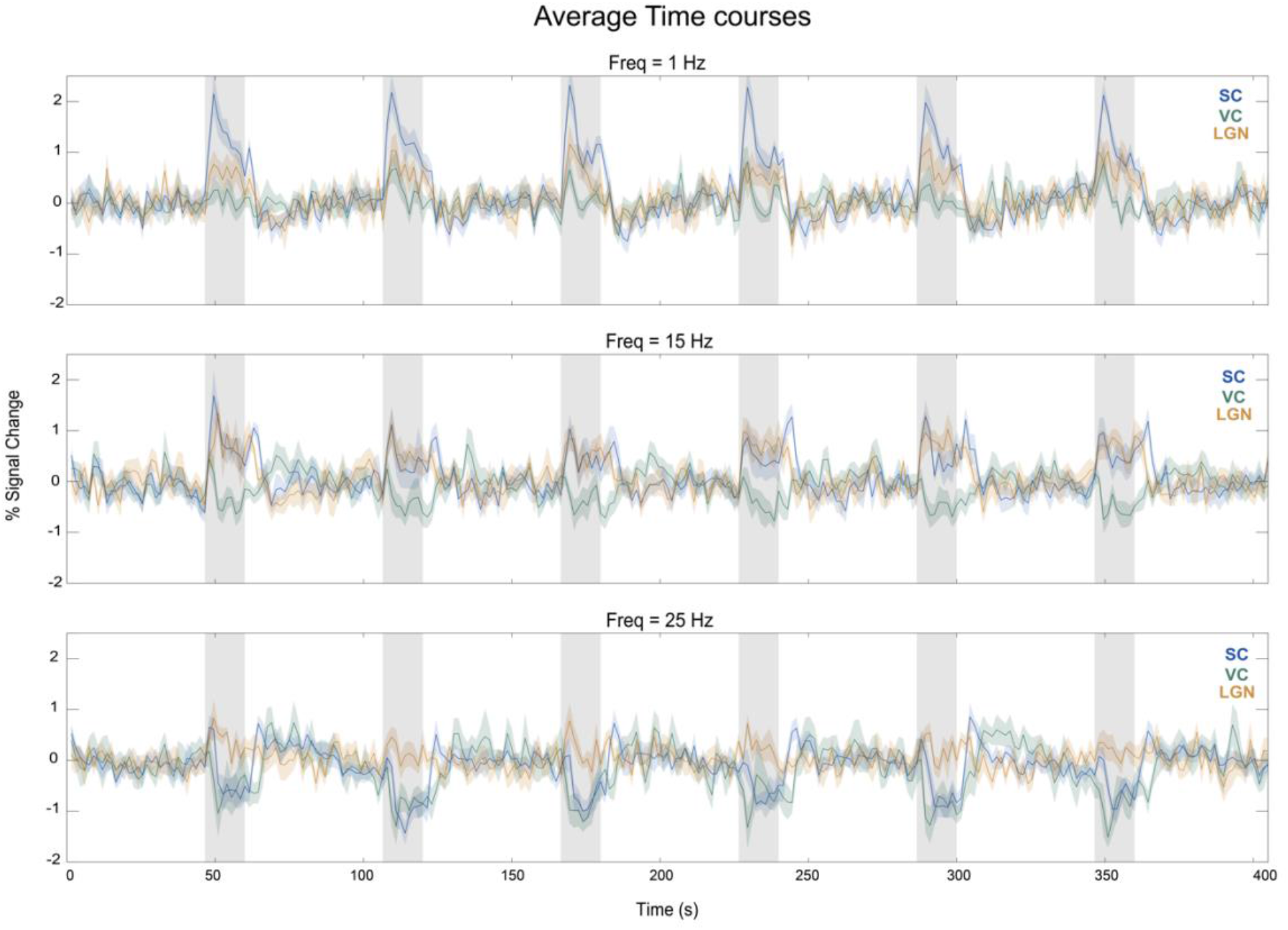
Average time-courses for the different visual pathway structures. Time-courses are shown for the three ROIs – VC (green), SC (blue) and LGN (orange) – for three different stimulation frequencies – 1 Hz, 15 Hz and 25 Hz. Results show robust fMRI signal changes upon visual stimulation and no evidence of habituation along the stimulation cycles.

Responses are consistent along runs and no habituation is observable along the cycles for any visual pathway structure.

##### fMRI Raw Data movie

Movie of raw data (4 fMRI acquisition slices) where the overall measured SNR is above 115. Data was acquired using a 9.4T BioSpec scanner (Bruker, Karlsruhe, Germany) with an 86mm quadrature resonator for transmittance and a 4-element array cryoprobe for signal reception. A SE-EPI sequence was used: TE/TR=40/1500 ms, partial Fourier coefficient=1.5, FOV=18×16.1 mm^2^, resolution=269×268 μm^2^, slice thickness=1.5 mm, t_acq_=7 min 30 s.

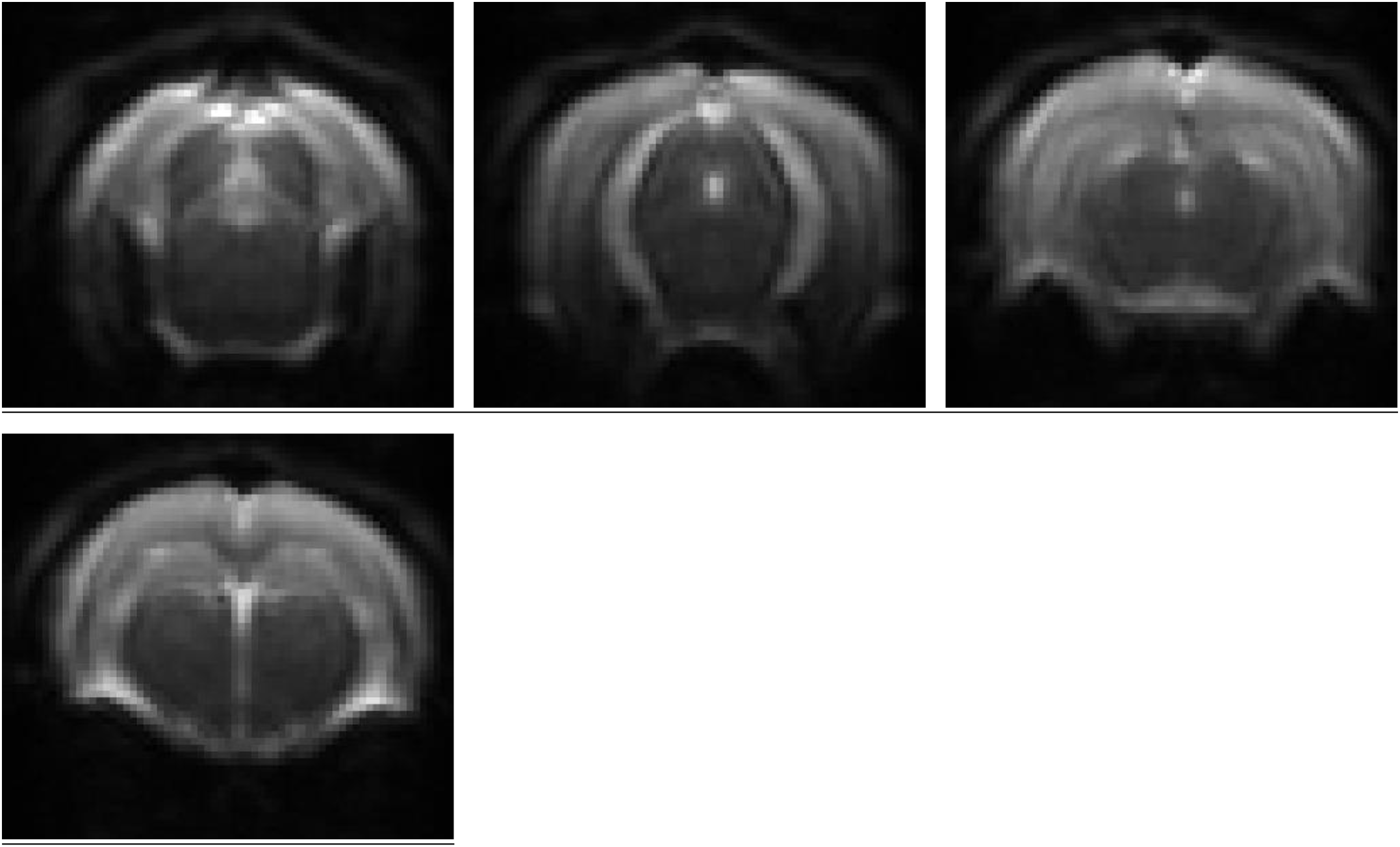

#### Electrophysiology

##### Zoomed in LFP signal amplitudes

Median LFP signal fluctuations reveal for the 1 Hz stimulation condition strong LFP oscillations induced by each flash. Closer inspection of the time profiles around the beginning and end of the stimulation period (**Figure S6**) reveals, similarly to the 1 Hz condition, that a single flash induces LFPs during the 15 Hz and 25 Hz stimulation period, albeit with smaller amplitude as the frequency increases. This feature is completely lost in the true continuous light stimulation condition. One similarity between the three conditions is the LFP onset oscillation while the offset oscillation is only present for the highest frequencies.

**Figure S6:**
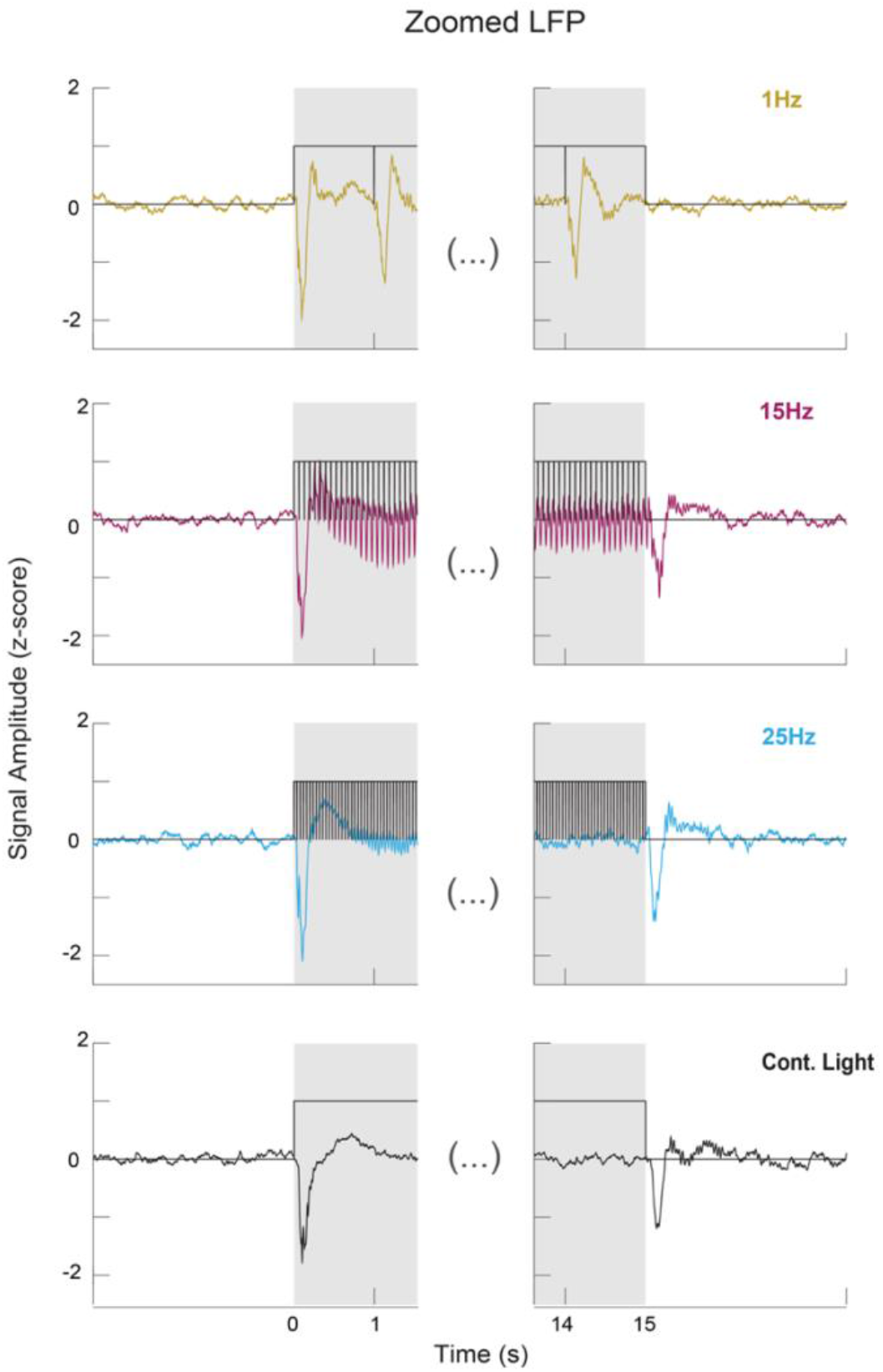
Zoomed mean LFP traces. Animal averaged zoomed traces for the beginning and end of stimulation for the 1, 15, 25 Hz and continuous light stimulation conditions. Black lines represent individual flashes. These plots confirm the individual flash induced oscillations and the absence of offset oscillations for the 1 Hz condition. For the 15Hz and 25 Hz conditions smaller individual flash induced oscillation are present along with marked onset and offset oscillations. For the continuous light condition only onset and offset oscillation are observed.

##### Spectrogram for remaining tested frequencies

In **Figure S7** we show, similarly to what is shown in **Figure 3C**, spectrograms between 0-50 Hz for the remaining tested frequencies.

**Figure S7:**
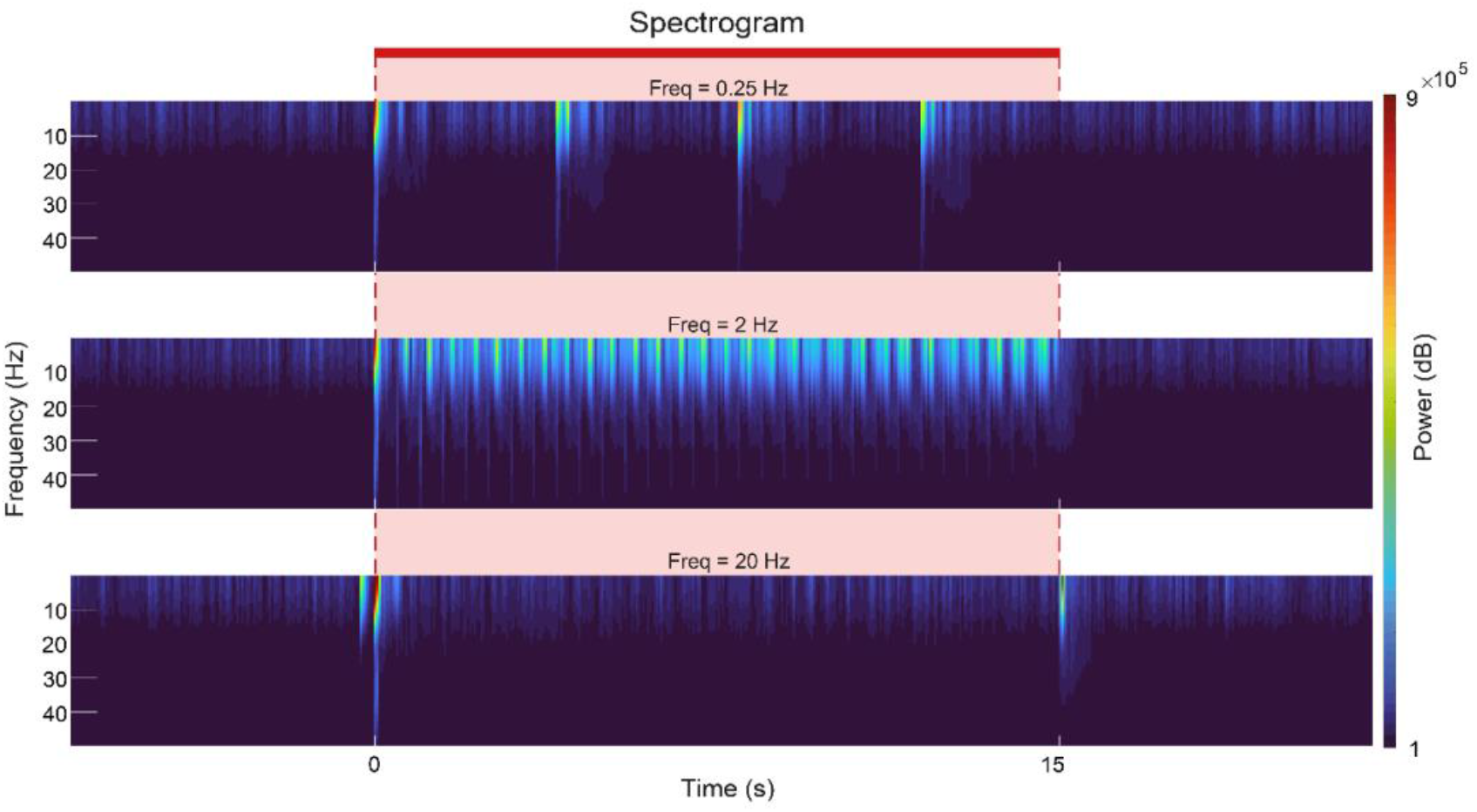
Spectrograms. **(A) Spectrograms between 1-50 Hz.** These plots confirm the individual flash induced power increases and the absence of offset oscillations for the 0.25 Hz and 2 Hz condition. For the 20 Hz condition onset and offset power increases are observed similarly to the 25 Hz;

##### fMRI and electrophysiology signals for a larger frequency space

To better characterise the evolution of electrophysiological signal along different frequencies a wider range of frequencies was tested and the extra frequencies are shown in **Figure S8**.

**Figure S8:**
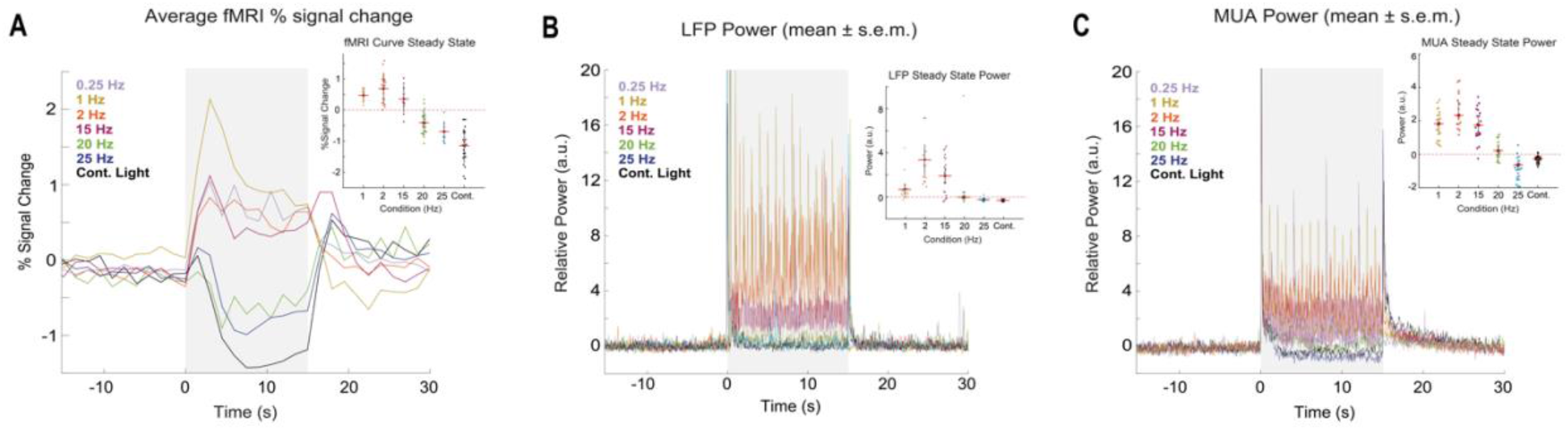
**(A) fMRI time profiles.** Higher stimulation frequencies lead to stronger SC NBRs; **(B) LFP relative power.** LFP power for all tested frequencies where stronger power reduction is observed for the 20, 25Hz and continuous light conditions; **(C) MUA relative power.** Similar trends as the ones observed for the LFP band. Interestingly high frequencies induced even a stronger MUA power reduction below baseline levels.

##### LFP power and fMRI percent signal change at Steady-state” Correlation

**Figure S9** shows the correlation between the LFP and fMRI signals during “steady-state”. Coefficients of ρ_Pearson_=0.84 (P=0.04, two sampled *t*-test) for the steady-state were achieved.

**Figure S9:**
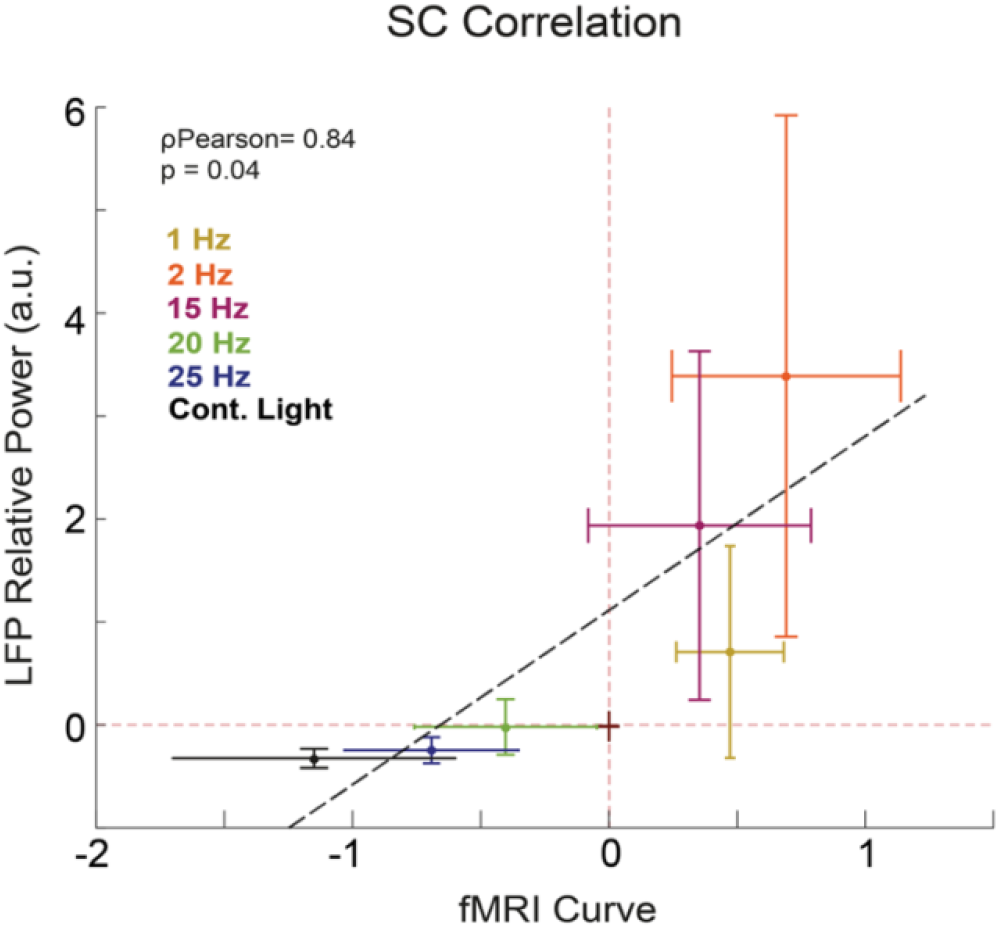
Correlation between fMRI signal and LFP power at steady-state. The correlation failed to be statistically significant with a coefficient of ρ_Pearson_=0.84 (P=0.04, two sampled *t*-test). NBRs at high stimulation frequencies correlate with strong LFP power reductions close to baseline levels. The circles represent the mean LFP power and fMRI percent signal change across animals while the error bars represent the standard deviation of the mean LFP power and fMRI percent signal change across runs.

##### LFP and MUA Convolution with HRF

In **Figure S10** the convolution of a typical HRF (peaking at 1 s) with electrophysiological power plots was performed in order to observe how such convolved signals would resemble fast fMRI responses. While onset and offset peaks between fast fMRI responses (N=6) and convolved signals appear well aligned, the amplitude of the negative fMRI responses observed at high stimulation frequencies cannot be completely explained. This means that others phenomena captured by the fMRI signals and not by the electrophysiological ones emphasize the negative amplitudes.

**Figure S10:**
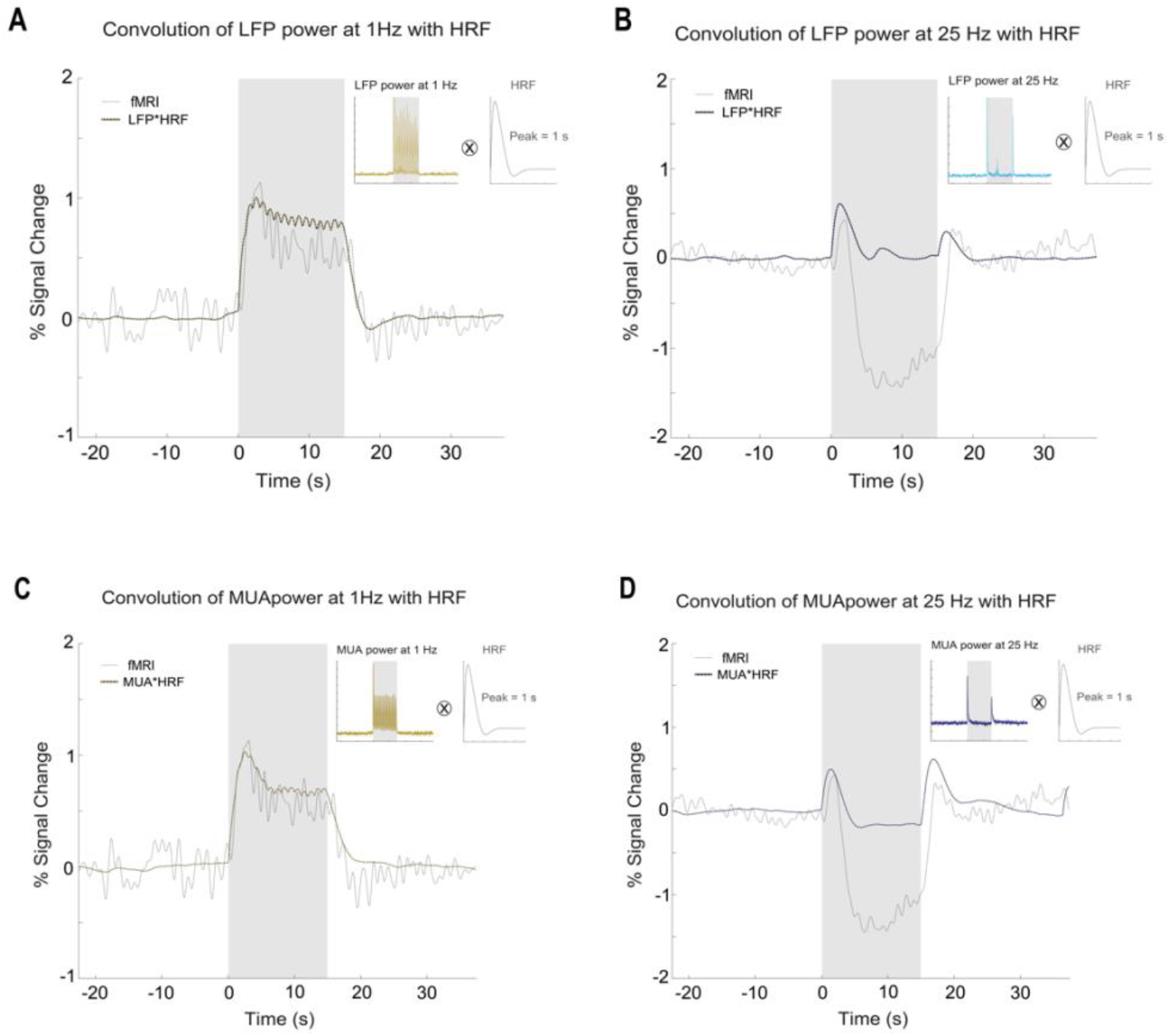
Convolution electrophysiological data with an HRF peaking at 1 sec. **LFP convolutions for the 1 Hz (A) and 25 Hz (B) condition.** The resulting convolved LFPs were compared with a fast fMRI acquisition (TR = 500 ms). Onset/offset peaks between the two curves appear aligned. **MUA convolution for the 1 Hz (D) and 25 Hz (E) condition.** The resulting convolved MUA was compared with a fast fMRI acquisition (TR = 500 ms). Onset/offset peaks between the two curves are aligned with onsets occurring ∼1.5-2sec after stimulation started and offsets peaking ∼1.8-2.3 sec after stimulation ended.

##### Ibotenic acid lesions

**Figure S11** depicts further results for the ibotenic acid lesions in V1 (N=10). As expected, V1 temporal profiles appear flat with no positive to negative fMRI signal shifts with increasing stimulation frequency as was observed for the control regime. Temporal profiles in LGN are very similar between the lesion and control group, suggesting that the V1 lesion does not strongly affect processing in LGN. Lesions were separated from the fMRI experiments by one week to avoid inflammation and swelling artefacts in the images.

**Figure S11:**
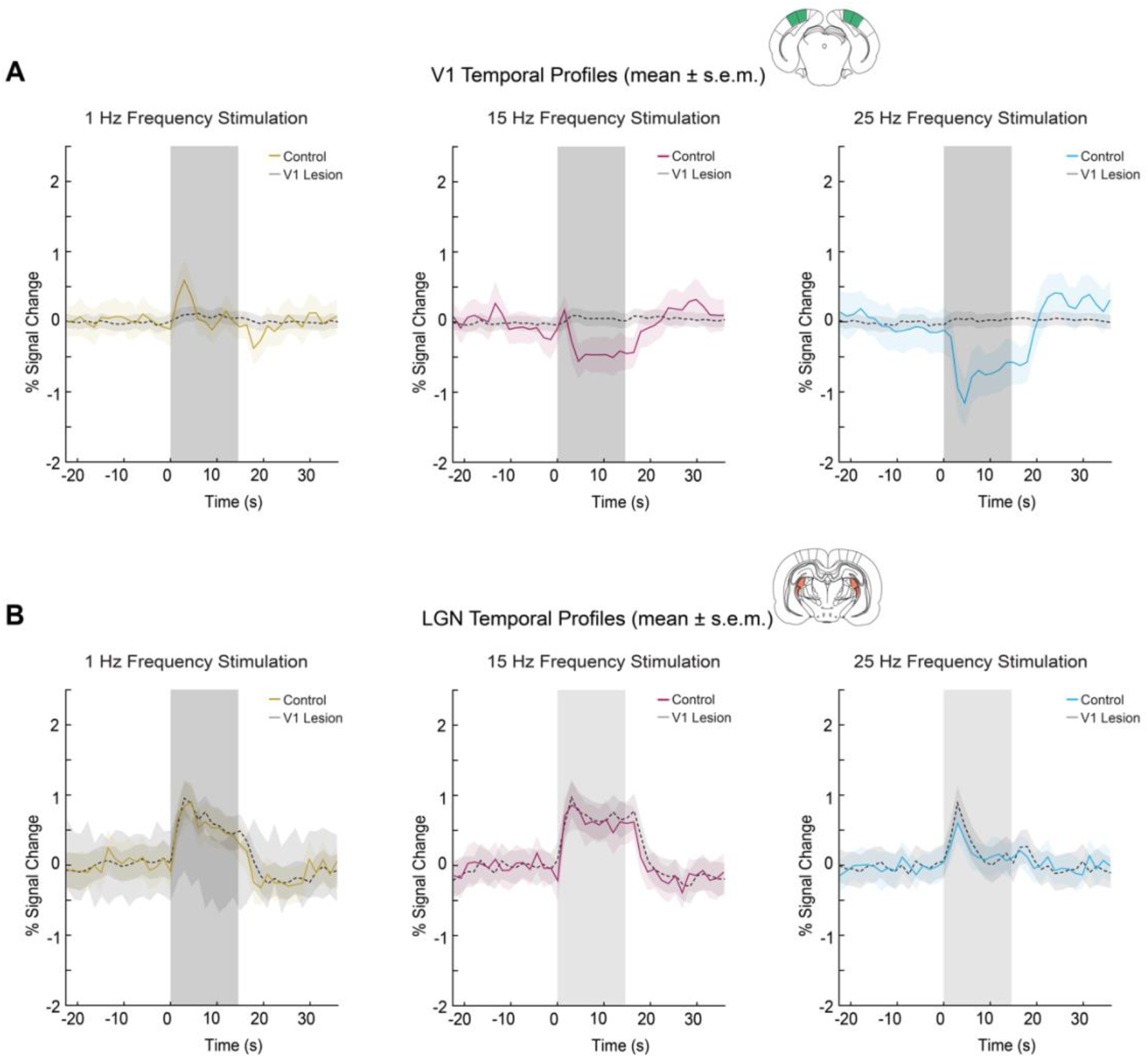
Cortical and thalamic fMRI temporal profiles after V1 ibotenic lesion (mean ± s.e.m. across animals). V1 profiles appear flat as expected from localized ibotenic acid lesions while LGN profiles appear similar to the control conditions with clear modulation as stimulation frequency increase but never reaching negative values.

## References

1. Nassi, J. J. & Callaway, E. M. Parallel processing strategies of the primate visual system. Nature Reviews Neuroscience vol. 10 360–372 Preprint at https://doi.org/10.1038/nrn2619 (2009).

2. Schlag, J. & Schlag-Rey, M. Through the eye, slowly; Delays and localization errors in the visual system. Nat Rev Neurosci 3, 191–200 (2002).

3. Siegle, J. H. et al. Survey of spiking in the mouse visual system reveals functional hierarchy. Nature 592, 86–92 (2021).

4. Sabesan, R., Schmidt, B. P., Tuten, W. S. & Roorda, A. The elementary representation of spatial and color vision in the human retina. Sci Adv 2, (2016).

5. Teh, K. L., Sibille, J., Gehr, C. & Kremkow, J. Retinal waves align the concentric orientation map in mouse superior colliculus to the center of vision. https://www.science.org (2023).

6. Basso, M. A., Bickford, M. E. & Cang, J. Unraveling circuits of visual perception and cognition through the superior colliculus. Neuron vol. 109 918–937 Preprint at https://doi.org/10.1016/j.neuron.2021.01.013 (2021).

7. White, B. J. et al. Superior colliculus neurons encode a visual saliency map during free viewing of natural dynamic video. Nat Commun 8, (2017).

8. Ge, X. et al. Retinal waves prime visual motion detection by simulating future optic flow. Science (1979) 373, (2021).

9. Li, Y. tang, Turan, Z. & Meister, M. Functional Architecture of Motion Direction in the Mouse Superior Colliculus. Current Biology 30, 3304–3315.e4 (2020).

10. de Malmazet, D., Kühn, N. K. & Farrow, K. Retinotopic Separation of Nasal and Temporal Motion Selectivity in the Mouse Superior Colliculus. Current Biology 28, 2961–2969.e4 (2018).

11. Beltramo, R. & Scanziani, M. A collicular visual cortex: Neocortical space for an ancient midbrain visual structure. https://www.science.org.

12. Feinberg, E. H. & Meister, M. Orientation columns in the mouse superior colliculus. Nature 519, 229–232 (2015).

13. Hafed, Z. M. & Chen, C. Y. Sharper, Stronger, Faster Upper Visual Field Representation in Primate Superior Colliculus. Current Biology 26, 1647–1658 (2016).

14. Arcaro, M. J., Honey, C. J., Mruczek, R. E., Kastner, S. & Hasson, U. Widespread correlation patterns of fMRI signal across visual cortex reflect eccentricity organization. doi:10.7554/eLife.03952.001.

15. Seabrook, T. A., Burbridge, T. J., Crair, M. C. & Huberman, A. D. Architecture, Function, and Assembly of the Mouse Visual System. (2017) doi:10.1146/annurev-neuro-071714.

16. Reber, M., Burrola, P. & Lemke, G. A relative signalling model for the formation of a topographic neural map. Nature 431, 847–853 (2004).

17. Eisen-Enosh, A., Farah, N., Burgansky-Eliash, Z., Polat, U. & Mandel, Y. Evaluation of Critical Flicker-Fusion Frequency Measurement Methods for the Investigation of Visual Temporal Resolution. Sci Rep 7, (2017).

18. Boström, J. E. et al. Ultra-Rapid vision in birds. PLoS One 11, (2016).

19. Yang, S. et al. The electroretinogram of mongolian gerbil (Meriones unguiculatus): Comparison to mouse. Neurosci Lett 589, 7–12 (2015).

20. Lisney, T. J., Ekesten, B., Tauson, R., Håstad, O. & Ödeen, A. Using electroretinograms to assess flicker fusion frequency in domestic hens Gallus gallus domesticus. Vision Res 62, 125–133 (2012).

21. Gilmour, G. S. et al. The electroretinogram (ERG) of a diurnal cone-rich laboratory rodent, the Nile grass rat (Arvicanthis niloticus). Vision Res 48, 2723–2731 (2008).

22. Schwartz, A. S. Electrophysiological Correlates of Flicker Perception in the Cat. Physiology and Behavior vol. 8 (1972).

23. Wells, E. F., Bernstein, G. M., Scott, B. W., Bennett, P. J. & Mendelson, J. R. Critical flicker frequency responses in visual cortex. Exp Brain Res 139, 106–110 (2001).

24. Schneider∼, C. W. Electrophysiological Analysis of the Mechanisms underlying the Critical Flicker Fusion Frequency. Vision Res. 1235–1244 (1968).

25. Lisney, T. J. et al. Behavioural assessment of flicker fusion frequency in chicken Gallus gallus domesticus. Vision Res 51, 1324–1332 (2011).

26. Mankowska, N. D. et al. Critical flicker fusion frequency: A narrative review. Medicina (Lithuania) vol. 57 Preprint at https://doi.org/10.3390/medicina57101096 (2021).

27. Landis’, C. Determinants of the Critical Flicker-Fusion Threshold. www.physiology.org/journal/physrev (1954).

28. Euler, T. & Wassle, H. Immunocytochemical Identification of Cone Bipolar Cells in the Rat Retina. THE JOURNAL OF COMPARATIVE NEUROLOGY vol. 361 (1995).

29. Umeton, D., Read, J. C. A. & Rowe, C. Unravelling the illusion of flicker fusion. Biol Lett 13, (2017).

30. Shumake, S. A., Smith, J. C. & Taylor, H. L. Critical Fusion Frequency in Rhesus Monkeys. Psychol Rec 18, 537–542 (1968).

31. Anderson, K. V, Keith, W. O. & Altered, S. Altered Response Latencies on Visual Discrimination Tasks in Rats with Damaged Retinas. Physiology and Behavior vol. 12 (1974).

32. Coile, D. C., et al. Behavioral Determination of Critical Flicker Fusion in Dogs. Phystology& Behavtor vol. 45.

33. Schwartz, A. S. & Cheney, C. Neural Mechanisms involved in the critical flicker frequency of the cat. Brain Reseach 1, 369–380 (1966).

34. Hendricks, J. Flicker Thresholds as Determined by a Modified Conditioned Suppression Procedure. Journal of the experimental analysis of behavior 9, (1966).

35. Rubene, D., Håstad, O., Tauson, R., Wall, H. & Ödeen, A. The presence of UV wavelengths improves the temporal resolution of the avian visual system. Journal of Experimental Biology 213, 3357–3363 (2010).

36. Nomura, Y. et al. Evaluation of critical flicker-fusion frequency measurement methods using a touchscreen-based visual temporal discrimination task in the behaving mouse. Neurosci Res 148, 28– 33 (2019).

37. Logothetis, N. K., Pauls, J., Augath, M., Trinath, T. & Oeltermann, A. Neurophysiological investigation of the basis of the fMRI signal. www.nature.com (2001).

38. Logothetis, N. K. What we can do and what we cannot do with fMRI. Nature 453, 869–878 (2008).

39. Goense, J., Merkle, H. & Logothetis, N. K. High-Resolution fMRI Reveals Laminar Differences in Neurovascular Coupling between Positive and Negative BOLD Responses. Neuron 76, 629–639 (2012).

## Supplementary References

40. Attwell, D. et al. Glial and neuronal control of brain blood flow. Nature vol. 468 232–243 Preprint at https://doi.org/10.1038/nature09613 (2010).

41. Fukuda, M., Poplawsky, A. J. & Kim, S. G. Time-dependent spatial specificity of high-resolution fMRI: Insights into mesoscopic neurovascular coupling: Spatial Specificity of fMRI. Philosophical Transactions of the Royal Society B: Biological Sciences vol. 376 Preprint at https://doi.org/10.1098/rstb.2019.0623 (2021).

42. Chen, C. Y. & Hafed, Z. M. Orientation and Contrast Tuning Properties and Temporal Flicker Fusion Characteristics of Primate Superior Colliculus Neurons. Front Neural Circuits 12, (2018).

43. Loop, M. S., Pekjchowski∼, S. & Snnm, D. C. Flicker Fusion in normal and binocularly deprived cats. Vision Res 20, 49–57 (1980).

44. Norton∼, A. C. & Clark, G. Effects of cortical and collicular lesions on brightness and flicker discriminations in the cat. Vision Res 3, 29–44 (1963).

45. Eberhard Dodt, B. Differentiation Between Rods and Cones by Flicker Electroretiriograpliy in Pigeon and Guinea Pig.

46. Simonson, E. & Brozek, J. Flicker Fusion Frequency Backgrou nd and Applications. www.physiology.org/journal/physrev (1952).

47. A Van De Grind, B. W., Grusser, O. & Lunkenheimer, H. Temporal Transfer Properties of the Afferent Visual System Psychophysical, Neurophysiological and Theoretical Investigations.

48. Schwartz, A. S. & Clark, G. Discrimination of intermittent photic stimulation in the rat without its striate cortex. (1956).

49. Grubb, M. S. & Thompson, I. D. Quantitative Characterization of Visual Response Properties in the Mouse Dorsal Lateral Geniculate Nucleus. J Neurophysiol 90, 3594–3607 (2003).

50. Hawken, M. J., Shapley, R. M. & Grosof, D. H. Temporal-Frequency selectivity in the monkey visual cortex. Vis Neurosci 13, 477–492 (1996).

51. Ann Williams, R., et al. Flicker Detection in the Albino Rat Following Light-induced Retinal Damage. Physiology & Behavior vol. 34.

52. Taravellaand, C. L. & Clark, G. Discrimination of Intermittent Photic Stimulation in Normal and Brain-Damaged Cats. EXPERIMENTAL NEUROLOGY vol. 7 (1963).

53. Mayo, J. P. & Sommer, M. A. Neuronal adaptation caused by sequential visual stimulation in the frontal eye field. J Neurophysiol 100, 1923–1935 (2008).

54. Platt, B. & Withington, D. J. Response habituation in the superficial layers of the guinea-pig superior colliculus in vitro. Neuroscience Letters vol. 221 (1997).

55. Harutiunian-Kozak, B., Dec, K. & Dreher1, B. HABITUATION OF UNITARY RESPONSES IN THE SUPERIOR COLLICULUS OF THE CAT.

56. Vidyasagar, T. R. Pattern Adaptation in Cat Visual Cortex is a co-operative phenomenon. Neuroscience 36, 175–79 (1990).

57. Oyster, C. W. & Takahashi, E. S. Responses of Rabbit Superior Colliculus Neurons to Repeated Visual Stimuli. www.physiology.org/journal/jn.

58. Binns, K. E. & Salt, T. E. Excitatory amino acid receptors modulate habituation of the response to visual stimulation in the cat superior colliculus. Visual Neuroscience vol. 12 (1995).

59. Dyer, R. S. & Annau, Z. Flash Evoked Potentials from Rat Superior Colliculus I. Pharmacology Biochemistry & Behavior vol. 6.

60. Binns, K. E. & Salt, T. E. Different roles for GABA(A) and GABA(B) receptors in visual processing in the rat superior colliculus. Journal of Physiology 504, 629–639 (1997).

61. Cerri, D. H. et al. Distinct neurochemical influences on fMRI response polarity in the striatum. bioRvix doi:10.1101/2023.02.20.529283.

62. Vo, T. T. et al. Parvalbumin interneuron activity drives fast inhibition-induced vasoconstriction followed by slow substance P-mediated vasodilation. Proc Natl Acad Sci U S A 120, (2023).

63. Devor, A. et al. Suppressed neuronal activity and concurrent arteriolar vasoconstriction may explain negative blood oxygenation level-dependent signal. Journal of Neuroscience 27, 4452–4459 (2007).

64. Stefanovic, B., Warnking, J. M. & Pike, G. B. Hemodynamic and metabolic responses to neuronal inhibition. Neuroimage 22, 771–778 (2004).

65. Shmuel, A. et al. Sustained Negative BOLD, Blood Flow and Oxygen Consumption Response and Its Coupling to the Positive Response in the Human Brain. Neuron vol. 36 (2002).

66. Sten, S. et al. Neural inhibition can explain negative BOLD responses: A mechanistic modelling and fMRI study. Neuroimage 158, 219–231 (2017).

67. Pasley, B. N., Inglis, B. A. & Freeman, R. D. Analysis of oxygen metabolism implies a neural origin for the negative BOLD response in human visual cortex. NeuroImage vol. 36 269–276 Preprint at https://doi.org/10.1016/j.neuroimage.2006.09.015 (2007).

68. Boorman, L. et al. Negative blood oxygen level dependence in the rat: A model for investigating the role of suppression in neurovascular coupling. Journal of Neuroscience 30, 4285–4294 (2010).

69. Shmuel, A., Augath, M., Oeltermann, A. & Logothetis, N. K. Negative functional MRI response correlates with decreases in neuronal activity in monkey visual area V1. Nat Neurosci 9, 569–577 (2006).

70. Boillat, Y., Xin, L., van der Zwaag, W. & Gruetter, R. Metabolite concentration changes associated with positive and negative BOLD responses in the human visual cortex: A functional MRS study at 7 Tesla. Journal of Cerebral Blood Flow & Metabolism 1–13 (2019) doi:10.1177/0271678X19831022.

71. Northoff, G. et al. GABA concentrations in the human anterior cingulate cortex predict negative BOLD responses in fMRI. Nat Neurosci 10, 1515–1517 (2007).

72. Muthukumaraswamy, S. D., Edden, R. A. E., Jones, D. K., Swettenham, J. B. & Singh, K. D. Resting GABA concentration predicts peak gamma frequency and fMRI amplitude in response to visual stimulation in humans. PNAS 106, 8356–8361 (2009).

73. Baltes, C., Radzwill, N., Bosshard, S., Marek, D. & Rudin, M. Micro MRI of the mouse brain using a novel 400 MHz cryogenic quadrature RF probe. NMR Biomed 22, 834–842 (2009).

74. Watson, C. & P. G. The Rat Brain in stereotaxic coordinates. (2007).

75. Pai, S., Erlich, J. C., Kopec, C. & Brody, C. D. Minimal impairment in a rat model of duration discrimination following excitotoxic lesions of primary auditory and prefrontal cortices. Front Syst Neurosci (2011) doi:10.3389/fnsys.2011.00074.

76. Francisco Londoi, S., Duncan Luce, R. & Green, D. M. Handbook of Perception: Detection, Discrimination and Recognition. vol. II (Academic Press Inc., 1974).

77. Aleci, C. Chapter 4 Psychophysics for non-Psychophysicists. in Detection and Discrimination Threshold. in Measuring the Soul (ed. Les Ulis: EDP Sciences) 1720 (2021).

