## Supplementary Material for "Rat superior colliculus encodes the transition between static and dynamic vision modes"

One Sentence Summary: The rat superior colliculus plays a critical role in discriminating temporal frequency.

†Correspondence:
Dr. Noam Shemesh
Champalimaud Research, Champalimaud Centre for the Unknown
Av. Brasilia 1400-038, Lisbon, Portugal.

Phone number: +351 210 480 000 ext. #4467.

**The PDF file includes:**

- Supplementary Results and Figures
    - Behaviour
        - Behaviour movie
        - Behaviour Discussion
    - Functional MRI
        - Oxygenation Percentage influence on fMRI responses
        - Spin Echo vs. Gradient Echo fMRI acquisitions
        - Stimulation Parameters influence on measured fMRI responses
        - Average time-courses for different stimulation frequencies
        - fMRI Raw Data movie
    - Electrophysiology
        - Zoomed in LFP signal amplitudes
        - Spectrogram for remaining tested frequencies
        - fMRI and electrophysiology signals for a larger frequency space
        - LFP power and fMRI percent signal change at Steady-state" Correlation
        - LFP and MUA Convolution with HRF
        - Ibotenic acid lesions

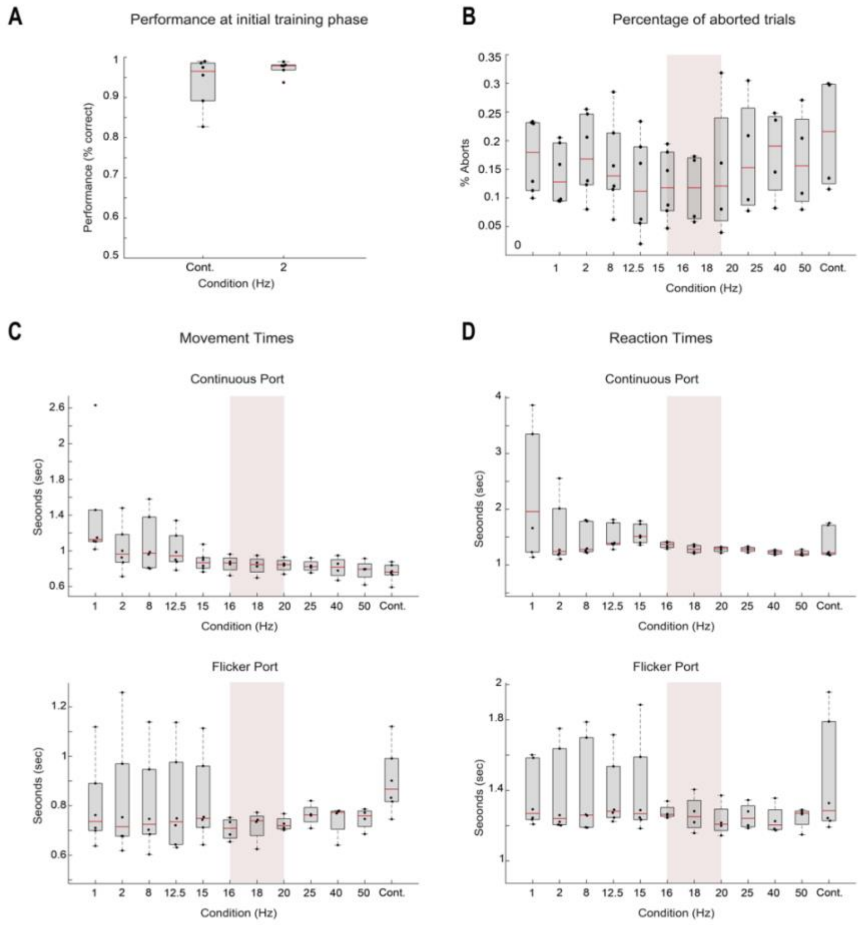

**Figure S1: Behaviour Results. (A) Performance at initial training phase.** Average performance of the animals for the initial training phase that included only the 2 Hz and "true continuous" stimuli. The values are shown for each of the presented frequencies and the mean is shown over animals with N=7; **(B) Percentage of aborted trials.** Percentage of trials aborted due to the animals attempting to respond before the mandatory minimum reaction time (1 second) reaching its end and the pure tone signalling the response period being played. The values are shown for each of the presented frequencies and the mean is shown over animals with N=7; **(C) Movement times**. Average over animals (N=7) of the movement times - time it takes the animal to reach the response port once it leaves the central port after the 1 second minimum-reaction time has elapsed - registered for the different frequencies presented. These are organized according to the report port for the trial, continuous port (top) and flickering port. The red-shaded area marks the calculated FFF ±2 Hz. **(D) Reaction times**. Average over animals (N=7) of the reaction times - time the animal is exposed to the stimulus, 1 second mandatory time plus the time the animal choses to linger until ready to respond - registered for the different frequencies presented. These are organized according to the report port for the trial, continuous port (**top**) and flickering port (**bottom**). The red-shaded area marks the calculated FFF threshold: 18±2 Hz.

### Functional MRI

<u>Oxygenation Percentage influence on fMRI responses</u>

Different oxygenation levels (medical air - 21% $O_2$; oxygen enriched air - 28% $O_2$; and hyperoxia - 95% $O_2$) were tested for two stimulation frequencies (2 Hz and 15 Hz) to investigate the sensitivity to measured positive and negative fMRI responses (**Figure S2**). Images were acquired with a SE-EPI sequence (TE/TR=43/1500 ms; FOV=18x16.1 mm; FA= 62°, resolution = 269x268 $\mu m^2$; slice thickness=1.5 mm). Results reveal that, for both frequency regimes, the hyperoxia oxygenation state (95% $O_2$) is the best regime to maximize fMRI responses, both positive and negative, along the rat visual pathway.

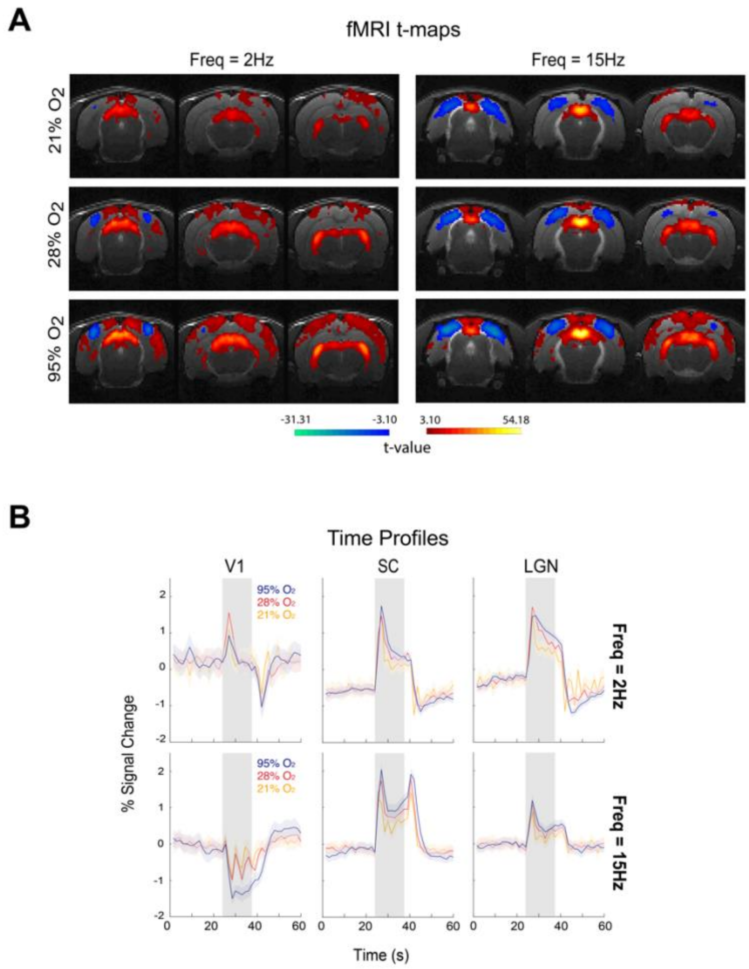


**Figure S2: fMRI signal modulation with different percentages of oxygen. (A) fMRI t-maps** for 23, 28 and 95% $O_2$ (p-value=0.001, cluster size=20 voxels). Bilateral fMRI responses along the main structures of the visual pathway are observed for both stimulation frequencies. Negative cortical fMRI responses along with positive subcortical fMRI responses are emphasized as the percentage of oxygen increases. For the 95% $O_2$ regime (hyperoxia) negative fMRI responses are observed in the V1M sub-regions (the monocular sub-region of the primary visual cortex) surrounded by positive responses in the rest of the primary visual cortex and secondary visual cortices. (**B) fMRI temporal profiles** for the different structures. An amplification of both positive and negative fMRI responses is confirmed for the hyperoxia condition. The amplification is mostly observed in cortical regions and more modest in subcortical regions. Legend: 21% $O_2$ (medical air)– yellow; 28% $O_2$ (oxygen enriched air) – orange; 95% $O_2$ (hyperoxia) – blue.

### Spin Echo vs. Gradient Echo fMRI acquisitions

To investigate if fMRI signal modulations observed at different stimulation frequency were specific to the chosen sequence, spin echo (SE) EPI, we tested several stimulation conditions with a gradient echo (GE) EPI sequence with similar parameters (TE/TR=15/1500 ms; FOV=18x16.1 mm; FA= 62º, resolution = 269x268 $\mu m^2$; slice thickness=1.5 mm) as in the SE-EPI sequence.

Results shown in **Figure S3** reveal similar signal modulations observed in GE-EPI fMRI as the ones observed in SE-EPI acquisitions. Negative cortical responses appear first at 15 Hz stimulation frequency however, these appear more modest than the ones observed with

SE-EPI acquisitions due to GE-EPI specific artifacts in these regions. SC responses show similar shifts from positive to negative responses as the stimulation frequency increases (at 25 Hz strong negative SC responses are observed). Due to enhanced specificity, the SE-EPI sequence was chosen for this study.

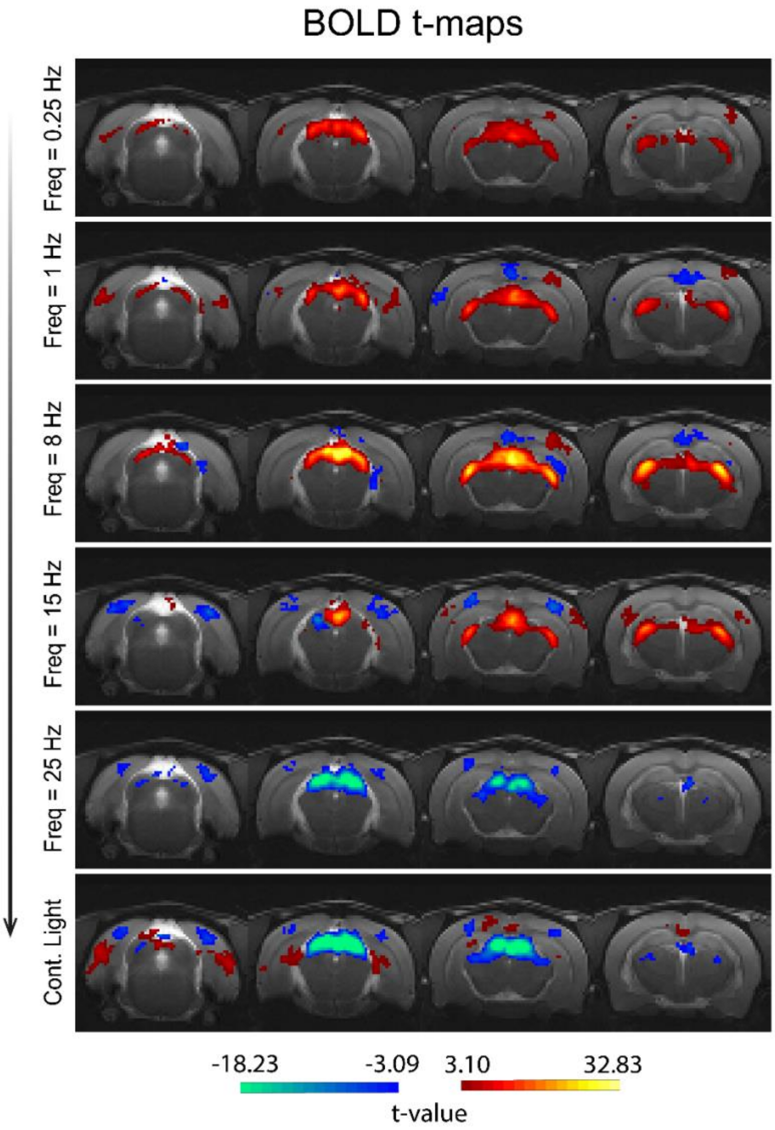


**Figure S3: GE-EPI fMRI t-maps along different stimulation frequencies** (cluster FDR corrected, p-value=0.001, cluster size=20 voxels). Bilateral fMRI responses are observed at the main visual pathway structures: SC, LGN and VC. Positive cortical responses at low stimulation frequencies are barely noticeable while stronger negative responses at frequencies higher than 15 Hz can be observed as in SE-EPI acquisitions. Subcortical responses also present signal modulations with strong negative responses in SC for frequencies higher than 25 Hz and reduced. LGN responses also appear modulated.

<u>Stimulation Parameters influence on measured fMRI responses</u>

From previous results we know that different frequencies with a constant flash duration of 10 ms (and consequently different ISIs) induce fMRI signal modulations. To further investigate the stimulation parameter (ISI vs flash duration) that has major influence in such signal modulations we tested fMRI signal modulations keeping the same ISIs but modulating the flash duration from 10 ms to 1 s (**Figure S4**). These acquisitions were performed using the chosen SE-EPI sequence: TE/TR=40/1500ms; FOV=18x16.1mm; resolution = 269x268μm$^2$; slice thickness=1.5mm.

Results show that different flash durations originate similar results revealing that the change in ISIs is the main factor inducing fMRI signal modulations.

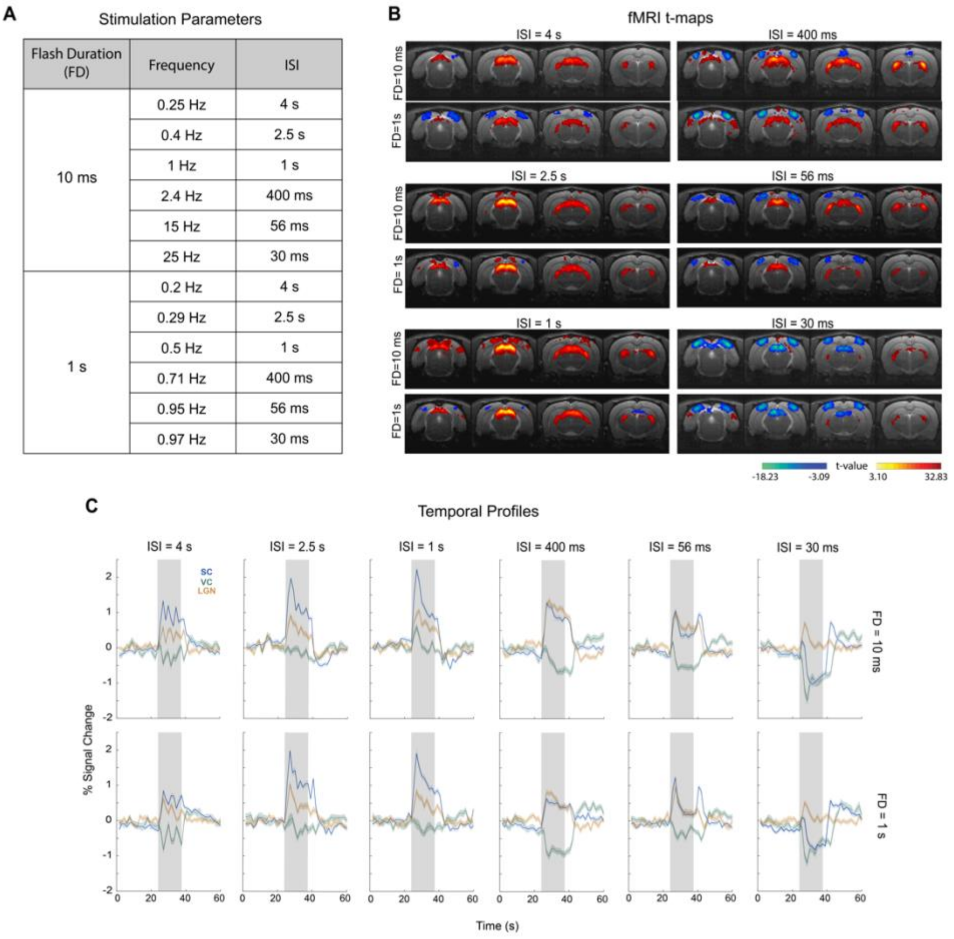


| Flash Duration (FD) | Frequency | ISI |
|---|---|---|
| 10 ms | 0.25 Hz | 4 s |
| | 0.4 Hz | 2.5 s |
| | 1 Hz | 1 s |
| | 2.4 Hz | 400 ms |
| | 15 Hz | 56 ms |
| | 25 Hz | 30 ms |
| 1 s | 0.2 Hz | 4 s |
| | 0.29 Hz | 2.5 s |
| | 0.5 Hz | 1 s |
| | 0.71 Hz | 400 ms |
| | 0.95 Hz | 56 ms |
| | 0.97 Hz | 30 ms |

**Figure S4: Effects of stimulation parameters on fMRI response modulations along the visual pathway. (A) Stimulation Parameters** for the different tested conditions. Two flash durations (FD) were tested, 10 ms and 1 s, while keeping similar ISIs. **(B) fMRI t-maps** (p-value=0.001, cluster size=8 voxels, FDR cluster correction). Maps appear similar for both flash durations along the tested ISIs with increased cortical and subcortical negative fMRI signals at high frequencies; **(C) fMRI Temporal Profiles.** To further investigate the how stimulation parameters affected fMRI signal modulations we plot the temporal profiles of signals with different FDs. These plots show similar curves for the three different ROIs (superior colliculus, SC, lateral geniculate thalamic nucleus, LGN, and visual cortex, VC) for the two different FDs. Modulations appear similar as the ISIs decrease. These results reveal that the ISI is the major factor inducing such changes.

Average time-courses for different stimulation frequencies

Time courses from the chosen SE-EPI sequence are shown in **Figure S5** for the different drawn ROIs along the visual pathway and different stimulation frequencies.

Responses are consistent along runs and no habituation is observable along the cycles for any visual pathway structure.

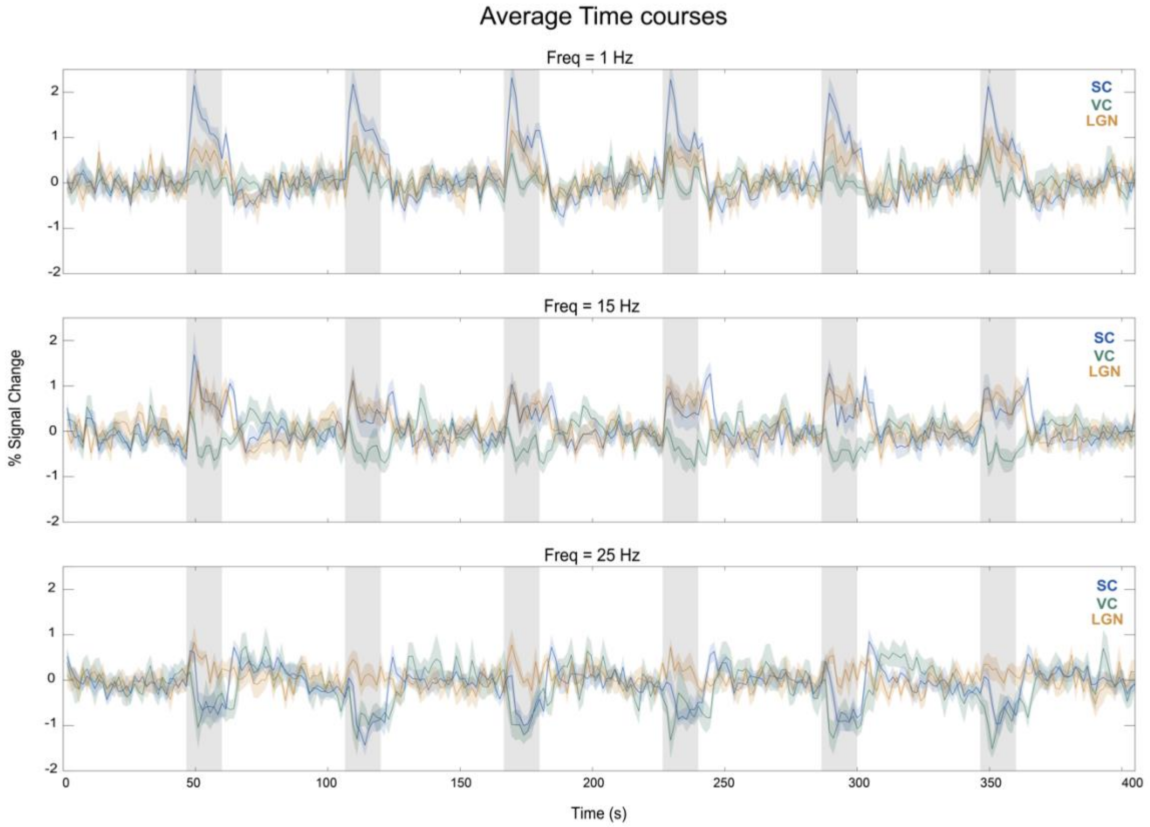


**Figure S5: Average time-courses for the different visual pathway structures.** Time-courses are shown for the three ROIs – VC (green), SC (blue) and LGN (orange) – for three different stimulation frequencies – 1 Hz, 15 Hz and 25 Hz. Results show robust fMRI signal changes upon visual stimulation and no evidence of habituation along the stimulation cycles.

fMRI Raw Data movie

Movie of raw data (4 fMRI acquisition slices) where the overall measured SNR is above 115. Data was acquired using a 9.4T BioSpec scanner (Bruker, Karlsruhe, Germany) with an 86mm quadrature resonator for transmittance and a 4-element array cryoprobe for signal reception. A SE-EPI sequence was used: TE/TR=40/1500 ms, partial Fourier coefficient=1.5, FOV=18x16.1 $mm^2$, resolution=269x268 $\mu m^2$, slice thickness=1.5 mm, $t_{acq}$=7 min 30 s.

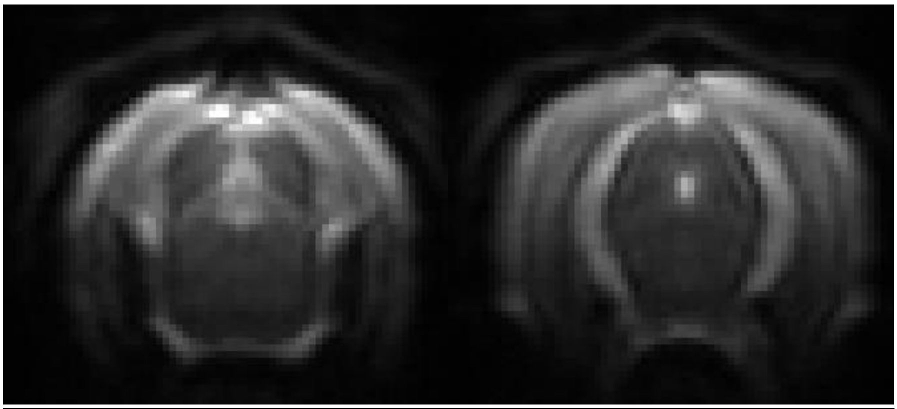

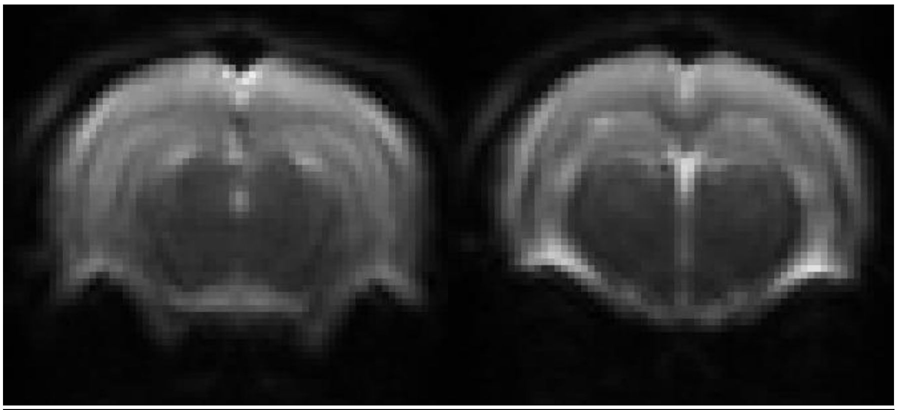

### Electrophysiology

Zoomed in LFP signal amplitudes

Median LFP signal fluctuations reveal for the 1 Hz stimulation condition strong LFP oscillations induced by each flash. Closer inspection of the time profiles around the beginning and end of the stimulation period (**Figure S6**) reveals, similarly to the 1 Hz condition, that a single flash induces LFPs during the 15 Hz and 25 Hz stimulation period, albeit with smaller amplitude as the frequency increases. This feature is completely lost in the true continuous light stimulation condition. One similarity between the three conditions is the LFP onset oscillation while the offset oscillation is only present for the highest frequencies.

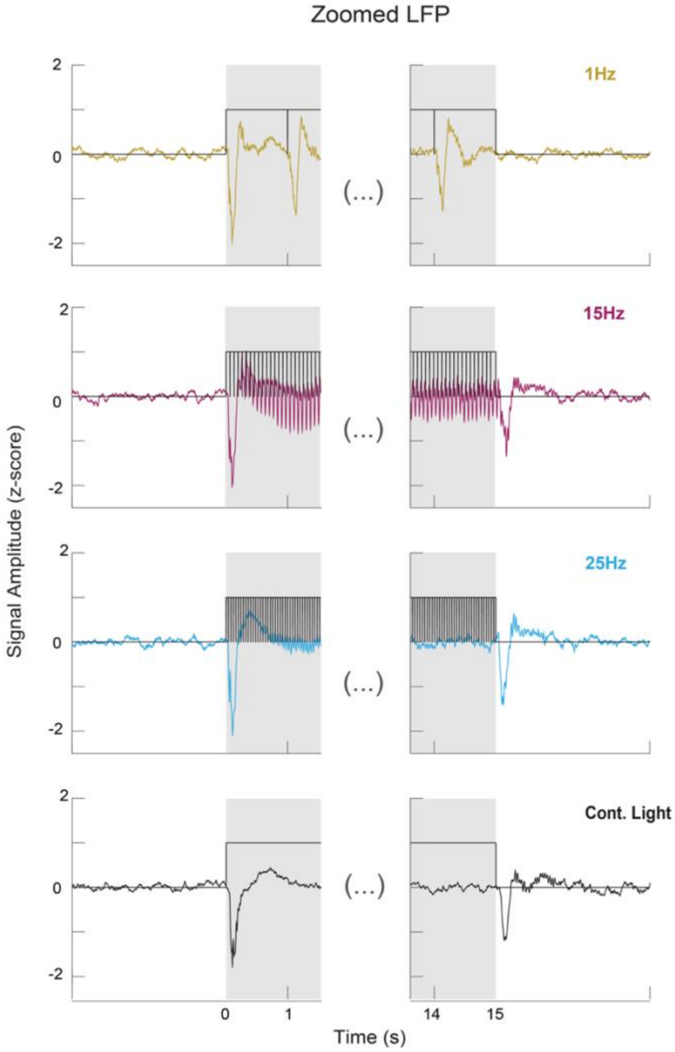


**Figure S6: Zoomed mean LFP traces.** Animal averaged zoomed traces for the beginning and end of stimulation for the 1, 15, 25 Hz and continuous light stimulation conditions. Black lines represent individual flashes. These plots confirm the individual flash induced oscillations and the absence of offset oscillations for the 1 Hz condition. For the 15Hz and 25 Hz conditions smaller individual flash induced oscillation are present along with marked onset and offset oscillations. For the continuous light condition only onset and offset oscillation are observed.

Spectrogram for remaining tested frequencies

In **Figure S7** we show, similarly to what is shown in **Figure 3C**, spectrograms between 0-50 Hz for the remaining tested frequencies.

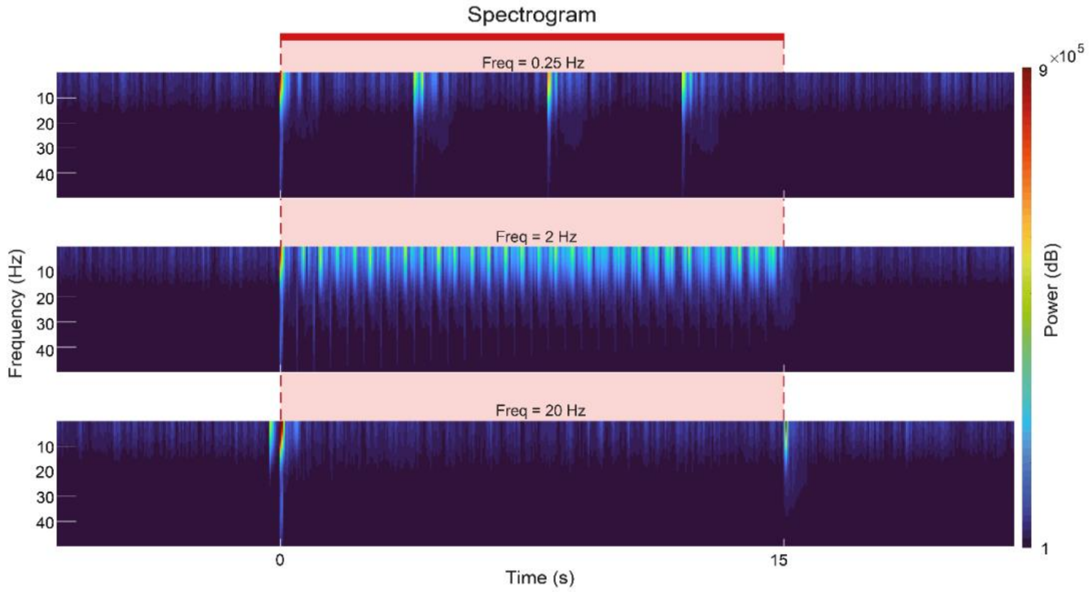


**Figure S7: Spectrograms. (A) Spectrograms between 1-50 Hz.** These plots confirm the individual flash induced power increases and the absence of offset oscillations for the 0.25 Hz and 2 Hz condition. For the 20 Hz condition onset and offset power increases are observed similarly to the 25 Hz;

fMRI and electrophysiology signals for a larger frequency space

To better characterise the evolution of electrophysiological signal along different frequencies a wider range of frequencies was tested and the extra frequencies are shown in **Figure S8**.

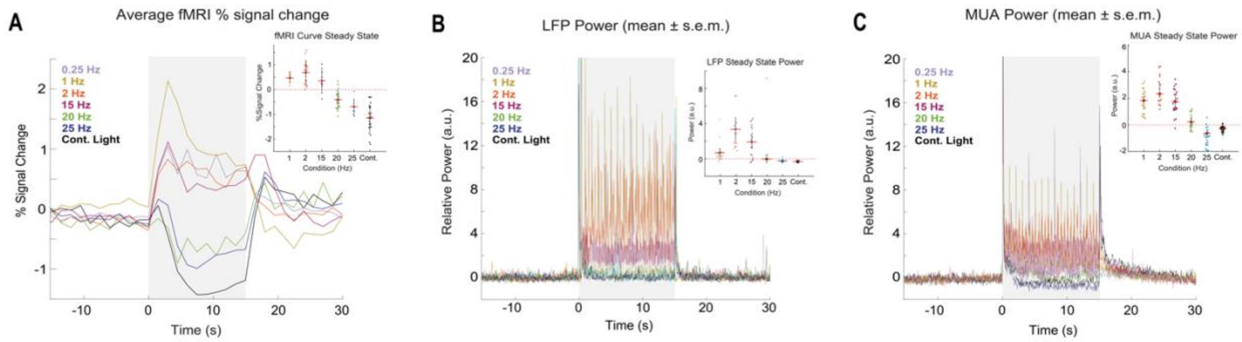


**Figure S8: (A) fMRI time profiles.** Higher stimulation frequencies lead to stronger SC NBRs; **(B) LFP relative power.** LFP power for all tested frequencies where stronger power reduction is observed for the 20, 25Hz and continuous light conditions; **(C) MUA relative power.** Similar trends as the ones observed for the LFP band. Interestingly high frequencies induced even a stronger MUA power reduction below baseline levels.

LFP power and fMRI percent signal change at Steady-state" Correlation

**Figure S9** shows the correlation between the LFP and fMRI signals during "steady-state". Coefficients of $\rho_{Pearson}$=0.84 (P=0.04, two sampled *t*-test) for the steady-state were achieved.

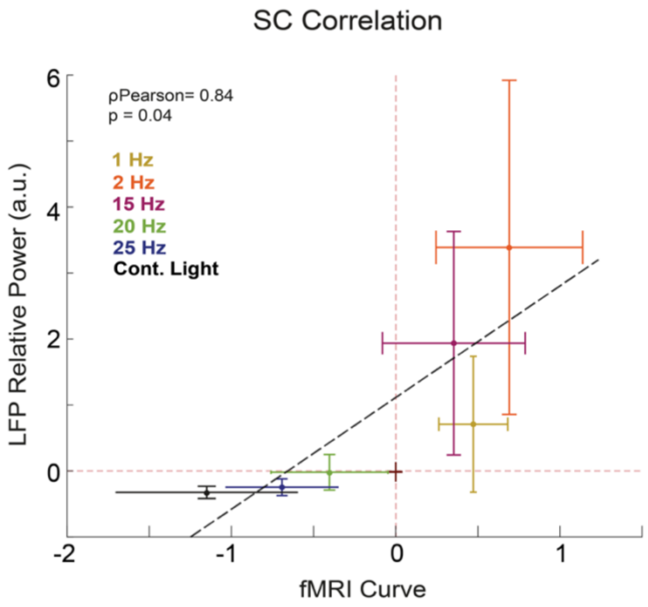


**Figure S9: Correlation between fMRI signal and LFP power at steady-state.** The correlation failed to be statistically significant with a coefficient of $\rho_{Pearson}$=0.84 (P=0.04, two sampled *t*-test). NBRs at high stimulation frequencies correlate with strong LFP power reductions close to baseline levels. The circles represent the mean LFP power and fMRI percent signal change across animals while the error bars represent the standard deviation of the mean LFP power and fMRI percent signal change across runs.

LFP and MUA Convolution with HRF

In **Figure S10** the convolution of a typical HRF (peaking at 1 s) with electrophysiological power plots was performed in order to observe how such convolved signals would resemble fast fMRI responses. While onset and offset peaks between fast fMRI responses (N=6) and convolved signals appear well aligned, the amplitude of the negative fMRI responses observed at high stimulation frequencies cannot be completely explained. This means that others phenomena captured by the fMRI signals and not by the electrophysiological ones emphasize the negative amplitudes.

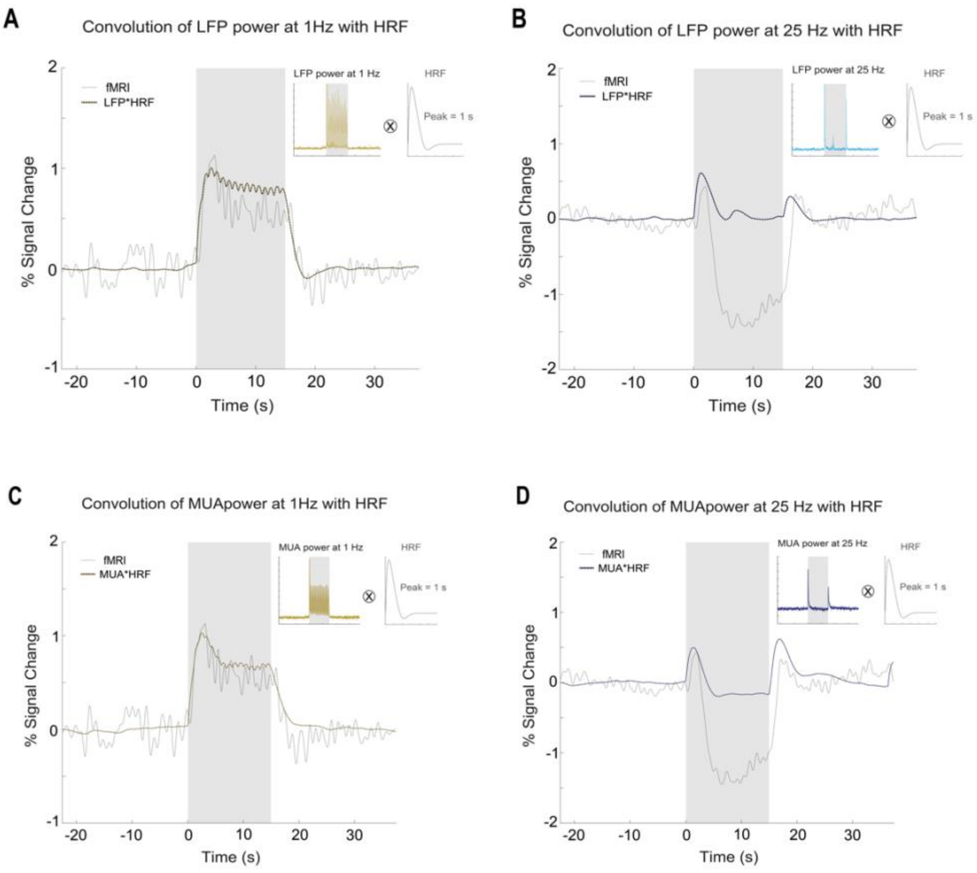


**Figure S10: Convolution electrophysiological data with an HRF peaking at 1 sec. LFP convolutions for the 1 Hz (A) and 25 Hz (B) condition.** The resulting convolved LFPs were compared with a fast fMRI acquisition (TR = 500 ms). Onset/offset peaks between the two curves appear aligned. **MUA convolution for the 1 Hz (D) and 25 Hz (E) condition.** The resulting convolved MUA was compared with a fast fMRI acquisition (TR = 500 ms). Onset/offset peaks between the two curves are aligned with onsets occurring ~1.5-2sec after stimulation started and offsets peaking ~1.8-2.3 sec after stimulation ended.

### Ibotenic acid lesions

**Figure S11** depicts further results for the ibotenic acid lesions in V1 (N=10). As expected, V1 temporal profiles appear flat with no positive to negative fMRI signal shifts with increasing stimulation frequency as was observed for the control regime. Temporal profiles in LGN are very similar between the lesion and control group, suggesting that the V1 lesion does not strongly affect processing in LGN. Lesions were separated from the fMRI experiments by one week to avoid inflammation and swelling artefacts in the images.

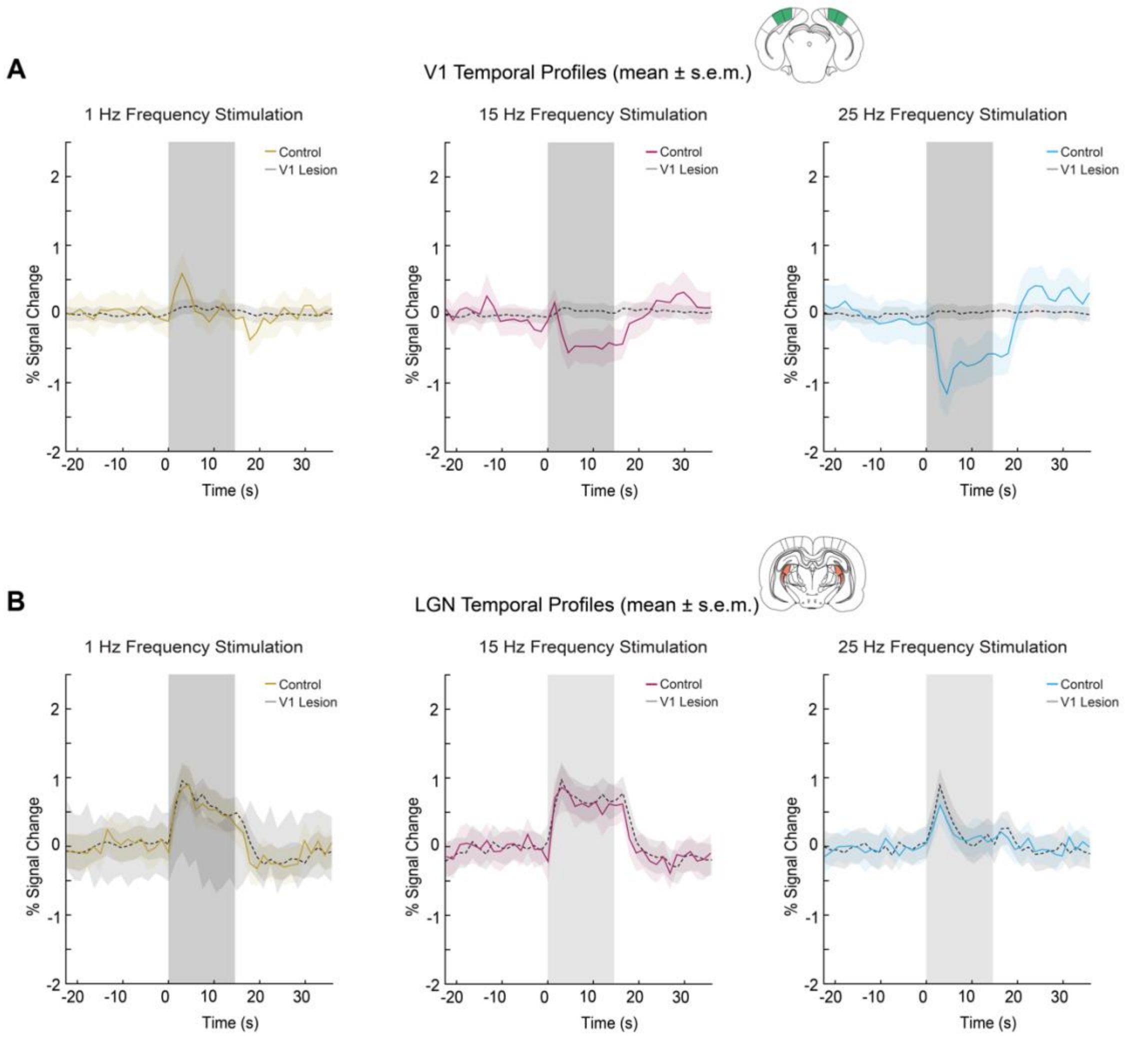


**Figure S11: Cortical and thalamic fMRI temporal profiles after V1 ibotenic lesion (mean ± s.e.m. across animals).** V1 profiles appear flat as expected from localized ibotenic acid lesions while LGN profiles appear similar to the control conditions with clear modulation as stimulation frequency increase but never reaching negative values.
